# Structural principles of peptide-centric Chimeric Antigen Receptor recognition guide therapeutic expansion

**DOI:** 10.1101/2023.05.24.542108

**Authors:** Yi Sun, Tyler J. Florio, Sagar Gupta, Michael C. Young, Quinlen F. Marshall, Samuel E. Garfinkle, Georgia F. Papadaki, Hau V. Truong, Emily Mycek, Peiyao Li, Alvin Farrel, Nicole L. Church, Shereen Jabar, Matthew D. Beasley, Ben R. Kiefel, Mark Yarmarkovich, Leena Mallik, John M. Maris, Nikolaos G. Sgourakis

## Abstract

Peptide-Centric Chimeric Antigen Receptors (PC-CARs), which recognize oncoprotein epitopes displayed by human leukocyte antigens (HLAs) on the cell surface, offer a promising strategy for targeted cancer therapy^1^. We have previously developed a PC-CAR targeting a neuroblastoma- associated PHOX2B peptide, leading to robust tumor cell lysis restricted by two common HLA allotypes^2^. Here, we determine the 2.1 Å structure of the PC-CAR:PHOX2B/HLA-A*24:02/β2m complex, which reveals the basis for antigen-specific recognition through interactions with CAR complementarity-determining regions (CDRs). The PC-CAR adopts a diagonal docking mode, where interactions with both conserved and polymorphic HLA framework residues permit recognition of multiple HLA allotypes from the A9 serological cross-reactivity group, covering a combined American population frequency of up to 25.2%. Comprehensive characterization using biochemical binding assays, molecular dynamics simulations, and structural and functional analyses demonstrate that high-affinity PC-CAR recognition of cross-reactive pHLAs necessitates the presentation of a specific peptide backbone, where subtle structural adaptations of the peptide are critical for high-affinity complex formation and CAR-T cell killing. Our results provide a molecular blueprint for engineering CARs with optimal recognition of tumor-associated antigens in the context of different HLAs, while minimizing cross-reactivity with self-epitopes.

## Introduction

Chimeric Antigen Receptor (CAR) T-cell therapy^3, 4^ has achieved remarkable efficacy for liquid tumors such as B-cell leukemias^5, 6^. On the other hand, while therapeutics for some solid tumors have seen anti-tumor activity^7–9^, they have been plagued by issues such as cytokine release syndrome and the lack of a sustained response. Thus, identifying tumor-specific antigens and developing highly selective CARs is critical for clinical applications^10^. While several promising cancer-specific antigens have been discovered, most proteins are expressed intracellularly and thus are undruggable by traditional CAR-T therapy^11, 12^. These inaccessible targets, however, are naturally degraded into peptides by the proteasome, which can be presented on the cell surface via major histocompatibility complex (MHC) class I molecules as peptide/MHC-I (pMHC) complexes, especially when aberrantly overexpressed in cancer cells^13^. Thus, CARs targeting endogenous peptides offer a new avenue for cancer immunotherapy^1^.

Neuroblastoma is an extracranial solid tumor responsible for 11% of pediatric cancer deaths^14^. With the completion of a phase 1-2 clinical trial, the success of a third-generation GD2-CAR T cell underscores the potential for CAR-T therapy for treating this malignancy^8^. To this end, we recently reported the development of a peptide-centric CAR (PC-CAR) 10LH targeting a neuroblastoma-enriched, unmutated peptide (QYNPIRTTF) derived from the PHOX2B oncoprotein^2^. While the single-chain fragment variable (scFv) based CAR displayed exquisite specificity for the PHOX2B antigen, its application was limited by HLA restriction to HLA-A*24:02 and HLA-A*23:01.

Here, we leverage insights from our 2.1 Å crystal structure of a 10LH:PHOX2B/HLA-A*24:02/β_2_ microglobulin (β_2_m) complex to expand the patient cohort amenable to CAR-T therapy. We characterize the molecular basis for peptide-centric recognition and explore the effect of HLA polymorphisms on 10LH binding. The PC-CAR 10LH can recognize the PHOX2B antigen presented by the HLA alleles from the A9 serological group, suggesting that it can be applied to treat up to a quarter of American, high-risk neuroblastoma patients. Through molecular dynamics (MD) simulations, biochemical assays, and *in situ* experiments, we report on the conformational plasticity of the peptide, enabling high-affinity complex formation and slow dissociation. Finally, we show that cytokine release by 10LH necessitates proper presentation of the R6 side chain and a conserved backbone, narrowing the cross-reactivity profile of the PC-CAR compared to naturally occurring TCR and TCR-like molecules. Our study uncovers structural principles required for tumor-associated HLA recognition, informing future PC-CAR T cell therapy development.

## Results

### The PC-CAR 10LH engages its PHOX2B/HLA-A*24:02 antigen through a prominent peptide selectivity filter

To understand the molecular basis of peptide-centric HLA recognition by 10LH, we determined the crystal structure of the 10LH:PHOX2B/HLA-A*24:02/β_2_m complex at a resolution of 2.1 Å (Fig. 1a, Extended Data Fig. 1, Extended Data Table 1, Supplementary Fig. 1a, and Supplementary Video 1). The PC-CAR engages the N- and C- termini of the MHC-I peptide binding groove via its complementarity-determining region 3 (CDR3) loops from the heavy and light chain, respectively (Fig. 1b). As reported in prior studies^15^, AlphaFold^16^ was unable to capture both the peptide interactions and the receptor docking orientation in its top 25 predictions of the 10LH:PHOX2B/HLA-A*24:02/β_2_m complex (Supplementary Video 2). We compared our 10LH structure to existing TCR:pHLA-I complexes and found that the CAR employed a parallel binding mode, with a relatively shallow docking angle (32.2°; Fig. 1c), which buried a moderate interface area on the peptide (Extended Data Fig. 2a). In contrast, the HLA/receptor interface was comparatively high relative to naturally occurring TCRs (Extended Data Fig. 2b), with interactions between the CDR loops and the α_1_ and α_2_ helices of HLA-A*24:02. HLA residues computationally identified as stabilizing the complex with 10LH were skewed towards one end of the MHC-I peptide-binding groove (Extended Data Fig. 2c), while TCRs displayed a more uniform footprint. Overall, the PC-CAR engages the PHOX2B/HLA-A*24:02/β_2_m complex in a manner resembling TCR:pHLA-I complexes, albeit with notable differences in its precise molecular interactions and extent of molecular surfaces, which justifies the formation of a high-affinity complex with a nanomolar equilibrium dissociation constant^2^, K_D_.

**Figure 1.**
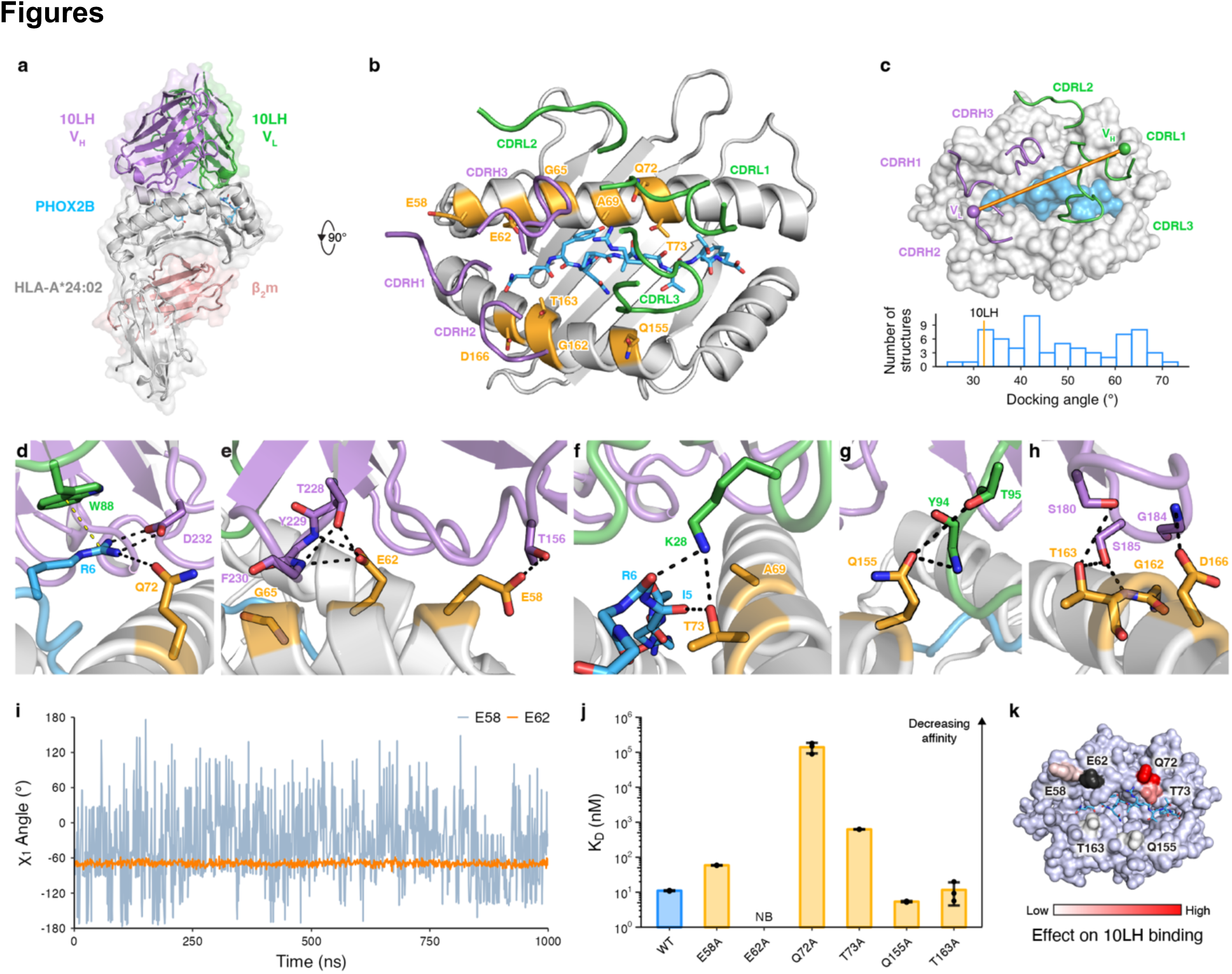
Structural basis for high-affinity 10LH:PHOX2B/HLA-A*24:02/β_2_m complex formation. **a,** Crystal structure of the 10LH:PHOX2B/HLA-A*24:02/β_2_m complex (PDB ID 8EK5). Surface representation is shown in lighter shades. The PHOX2B peptide is shown as sticks. **b,** Top-down view of the structure shown in **a**. The 10LH CDR loops are shown as cartoon tubes. The HLA framework residues (orange) and the PHOX2B peptide (blue) are shown in sticks. **c,** Top-down view of 10LH:PHOX2B/HLA-A*24:02/β_2_m shown in surface representation. The 10LH interdomain vector is drawn between the centroids of the conserved disulfide bonds in the heavy (purple) and light (green) chains. A histogram of TCR docking angles is calculated using the TCR interdomain vectors. The corresponding value for 10LH is indicated by an orange line at 32.2°. **d- h,** Close-up view of 10LH and PHOX2B/HLA-A*24:02/β_2_m interactions for the R6 in PHOX2B (**d**) and the framework residues along the α_1_ (**e**, **f**) and α_2_ (**g**, **h**) HLA helices. Hydrogen bonds and cation-π interactions are indicated by black and yellow dashed lines, respectively. **i,** The χ_1_ side chain torsion angle from MD simulations of the 10LH:PHOX2B/HLA-A*24:02 complex for E58 and E62. Data are mean across n = 3 independent 1 μs runs. **j**, SPR determined K_D_ values for 10LH:PHOX2B/HLA-A*24:02/β_2_m with alanine mutated framework residues. Data are mean ± SD for n = 3 technical replicates. K_D_, equilibrium dissociation constant; NB, no binding. **k,** The fold change of K_D_ for the alanine mutated 10LH:PHOX2B/HLA-A*24:02/β_2_m compared to the WT mapped on the HLA-A*24:02 surface. E62A (NB) is colored black.

We next focused on interactions in the PC-CAR:PHOX2B/HLA-A*24:02/β_2_m complex, which conferred antigen specificity. Q72 of HLA-A*24:02 interacts with R6 from PHOX2B, orienting the Arg side chain for salt bridge formation with D232 from CDRH3 (Fig. 1d). This network of contacts is stabilized further by a cation-π interaction with W88 from CDRL3, forming an exquisite peptide selectivity filter. Stable docking of the PC-CAR to the pHLA complex is mediated by multiple contacts with amino acids along the α_1_ and α_2_ helices, termed framework residues. On the α_1_ helix, E58 forms a hydrogen bond with T156 of 10LH CDRH1, and E62 forms three hydrogen bonds with CDRH3 residues (Fig. 1e). Moreover, K28 from the 10LH CDRL3 bridges both the peptide and HLA through a network of contacts with the R6 backbone carbonyl and the T73 side chain hydroxyl group (Fig. 1f). Four framework residues on the α_2_ helix form additional contacts with the CDRH2 and CDRL3 loops (Fig. 1g, h). To further investigate the stability of identified contacts between HLA-A*24:02 framework residues and 10LH, we conducted molecular dynamics (MD) simulations of the PC-CAR:pMHC platform at the 1 μs timescale, followed by experimental surface plasmon resonance (SPR) binding. Overall, our MD simulations showed a stable ternary complex, with relatively low mean root mean square fluctuation (RMSF) values compared to the crystallographic coordinates (Extended Data Fig. 3a). However, analysis of χ_1_ side chain torsion angles for the HLA framework residues identified in our structure revealed a range of dynamic effects (Extended Data Fig. 3b). For instance, while the E62 side chain is rigid along the simulation trajectory, E58 samples a wide range of χ_1_ angles suggesting a more transient interaction with the CDRH3 loop (Fig. 1i). Using a panel of HLA-A*24:02 point mutants, we observed a range of effects on 10LH binding by SPR (Fig. 1j, Extended Data Fig. 4, and Supplementary Fig. 2). Consistent with our MD simulations and analysis of the structure, E62A completely abrogated binding while E58A, Q155A, and T163A had a more subtle effect. Mapping these effects on the surface of HLA-A*24:02 revealed interaction hotspots (Fig. 1k) formed by E62 as well as Q72 and T73, residues that are adjacent to R6 of PHOX2B. Thus, these results demonstrate that high-affinity PC-CAR:PHOX2B/HLA-A*24:02/β_2_m complex formation emerges from the specific recognition of both R6 on PHOX2B, in addition to key framework residues on HLA-A*24:02. This is reminiscent of how TCRs engage their pMHC-I antigens^17, 18^, albeit with a strict requirement for a precise network of contacts formed by the Q72-R6-D232-W88 tetrad, which bridges elements from all three components of the molecular complex.

### 10LH interactions with polymorphic HLA framework residues enable recognition of multiple allotypes

Our structural analysis showed that a network of interactions with HLA framework residues was critical for 10LH complex formation. Thus, sequence divergence of HLA allotypes from the optimal HLA-A*24:02 interface is likely to disrupt antigen-specific recognition, despite maintaining PHOX2B presentation. We hypothesized that 10LH could recognize the PHOX2B peptide presented by members of the A9 serological cross-reactivity group (CREG)^19^, including multiple allotypes beyond the previously identified^2^ HLA-A*24:02 and HLA-A*23:01 (Fig. 2a). Upon prediction of PHOX2B binders among common (global freq. > 0.1%) HLAs by NetMHCPan 4.1^20^ (n = 219), we computed the sequence conservation for different framework residues (Fig. 2b). While Q72 is completely conserved across PHOX2B binding alleles, other positions including E62, G65, and A69 are highly polymorphic (Fig. 2c). These residues cluster on one face of the α_1_ helix, and form an epitope previously implicated in HLA alloreactivity^21^. Guided by the observed polymorphisms, we evaluated the effect of introducing specific HLA-A*24:02 substitutions on binding to 10LH by SPR (Fig. 2d, Extended Data Fig. 5, and Supplementary Fig. 3). Our results show that G65 substitutions to the arginine or glutamine polymorphism found in different HLAs abrogate binding, likely due to steric clashes with CDRH3. The E62Q and E62R substitution also resulted in a loss of binding, consistent with our previous alanine mutagenesis results. Substitution of A69 and G162 with bulkier, charged amino acids led to an increase of K_D_ values by two to three orders of magnitude, likely due to disruption of hydrophobic interactions. The D166E substitution had a more subtle effect. Thus, point mutations of polymorphisms found in PHOX2B-presenting alleles provide a strong basis for the HLA restriction of PHOX2B antigen recognition by 10LH.

**Figure 2.**
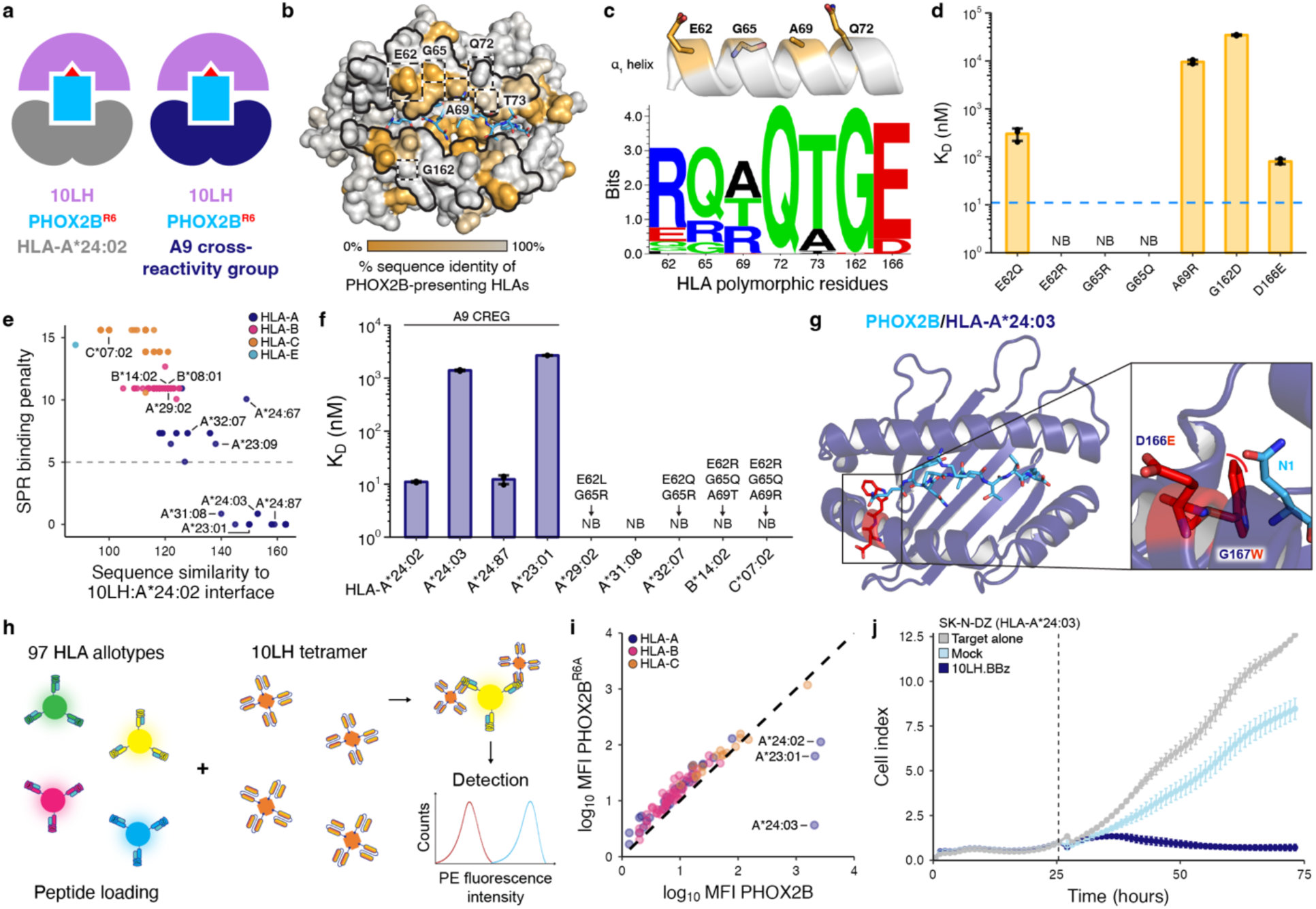
Key framework residues on HLA molecules confer allelic interactions with 10LH. **a,** Schematic depicting cross-HLA interactions by 10LH. **b,** Sequence identity across 219 HLA-I alleles computationally predicted to present the PHOX2B peptide mapped onto the HLA-A*24:02 surface (PDB ID 8EK5). The α_1_ and α_2_ helices and HLA framework residues are denoted by the black outlines and dashed boxes, respectively, on the HLA-A*24:02 surface. **c,** Schematic depicting the HLA framework residues on the α_1_ helix. Below, a sequence motif of the HLA framework residues across common alleles predicted to bind the PHOX2B peptide (n = 219). Created using Seq2Logo. **d,** SPR determined K_D_ values for PHOX2B/HLA-A*24:02/β_2_m with site-directed mutations of the framework residues. Data are mean ± SD for n = 3 technical replicates. The blue dashed line depicts the K_D_ of the WT PHOX2B/HLA-A*24:02/β_2_m to 10LH. **e,** Scatter plot of sequence similarity of HLA interface residues (BLOSUM62 score) against the SPR binding penalty for the predicted 10LH binders. The SPR binding penalty was calculated by adding the log base 10 ratio of the mean K_D_ value for each polymorphism in a given allele to the wild-type K_D_. The gray dashed line indicates the presence of at least one non-binding polymorphism. **f,** SPR determined K_D_ values for selected HLA allotypes. Polymorphisms among the framework residues are listed above non-binding alleles. Data are mean ± SD for n = 3 technical replicates. NB, no binding. **g,** Structural model of PHOX2B/HLA-A*24:03 with polymorphisms relative to HLA-A*24:02 denoted in red sticks. The zoom-in details the steric clash (red curve) between the polymorphic residue (red) and the PHOX2B peptide (blue). **h,** A summary of SAB screening, where beads are pre-incubated over weekend (o/w) with PHOX2B or PHOX2B^R6A^ peptides, and stained by 10LH PE-tetramers for a shift in MFI. **i,** Correlation of log_10_(MFI) for 10LH stained HLA allotypes pre-incubated with PHOX2B and PHOX2B^R6A^. Data are mean for n = 3 technical replicates with SD and individual data points shown in Extended Data Fig. 8. The black dashed line represents a conceptual 1:1 correlation (no difference between PHOX2B- and PHOX2B^R6A^- dependent 10LH interactions). **j,** *In-vitro* cytotoxic activity of 10LH.BBz CAR T cells against HLA-A*24:03 neuroblastoma cell line SK-N-DZ. 10LH.BBz CAR T cells were co-cultured (black dashed line indicates addition of effector cells to target cells) at a 3:1 E:T ratio with SK-N-DZ target cells. Target cell viability was determined using the cell index of the xCELLigence Real-Time Cell Analysis (RTCA) system. Target cell index was normalized to the timepoint immediately before addition of effector cells. Values represent mean ± SD using effector cells from n = 2 biological donors, in triplicates.

We used our measured K_D_ values for individual framework residue substitutions on HLA-A*24:02 to calculate a SPR binding penalty for all PHOX2B-presenting alleles and plotted these values against their BLOSUM62^22^ sequence similarity to HLA-A*24:02 with respect to interface residues in our 10LH complex structure as determined by PDBePISA^23^. We observed a partitioning of the allelic landscape into two distinct sets, with a clear cluster of HLA allotypes having a high sequence identity to HLA-A*24:02 and low binding penalty (Fig. 2e). We then conducted SPR binding experiments for a subset of HLAs from both sets (Fig. 2f, Extended Data Fig. 6, Supplementary Fig. 1b, and Supplementary Fig. 4), confirming interactions with 10LH for members of the A9 CREG, defined by the A23 and A24 serological splits. Notably, our identified 10LH binders included HLA-A*31:08, which was separate from the A9 cross-reactivity group. Analysis of this allotype’s sequence relative to HLA-A*24:02 revealed extensive polymorphisms in the HLA groove (Extended Data Fig. 7a), likely resulting in an altered conformation of the PHOX2B antigen and, therefore, impaired binding to 10LH as shown by our SPR data (Extended Data Fig. 7b). For alleles from the A9 CREG, an approximately 100-fold increase in K_D_ (micromolar range, as opposed to 11 nM for HLA-A*24:02) was observed. This could be attributed to the altered presentation of the PHOX2B antigen resulting from specific MHC-I groove polymorphisms as shown by our structural modeling for HLA-A*24:03 and HLA-A*23:01 (Fig. 2g and Extended Data Fig. 7c, d). Conversely, HLA-A*23:09 and HLA-A*24:67 from the A9 CREG were predicted to not bind to 10LH due to the deleterious G65R framework residue polymorphism. Other non-binding alleles outside of the A9 CREG, such as HLA-B*14:02, showed polymorphisms within multiple framework residues, leading to steric hindrance with the 10LH CDRs (Extended Data Fig. 7e, f). One selected allele, HLA-B*08:01, did not form stable complexes with the PHOX2B peptide despite the prediction by NetMHCPan 4.1^20^ (Supplementary Fig. 4). Thus, our structural modeling and biochemical analysis of framework residue polymorphisms identifies a set of PHOX2B-presenting HLA alleles that can cross-react with 10LH.

We further evaluated the cross-reactivity with the common HLA alleles using a single-antigen bead (SAB) assay^24^ (Fig. 2h). Inspection of mean fluorescence intensity (MFI) for the positive versus control experiments identified PHOX2B-dependent HLA interactions with 10LH for allotypes from the A9 CREG, in agreement with our SPR results (Fig. 2i and Extended Data Fig. 8). Using an exemplar, novel allele from this group, we demonstrated 10LH CAR-T cell cytotoxicity and cytokine release against an HLA-A*24:03 neuroblastoma cell line (Fig. 2j, Extended Data Fig. 9, and Supplementary Fig. 5). Overall, our results provide strong support that 10LH can bind to several additional allotypes from the A9 CREG. Thus, up to 25.2% of the American neuroblastoma patient cohort is amenable to 10LH CAR-T therapy, not considering linkage disequilibrium among HLA alleles (Extended Data Table 2).

### Peptide backbone plasticity enables high-affinity complex formation with 10LH

To examine the role of PHOX2B structural adaptations in specific recognition by 10LH, we compared the peptide conformations presented by HLA-A*24:02 in the apo and 10LH-bound states (Fig. 3a). A prominent backbone change, highlighted by a 1.7 Å displacement of the P4 carbonyl oxygen, enables an optimal presentation of the R6 side chain rotamer for interaction with the 10LH CDRs. To further characterize this conformational transition, we performed MD simulations of PHOX2B/HLA-A*24:02 in its apo and PC-CAR-bound states. Simulations starting from the coordinates of the apo structure sampled a bimodal distribution of Ψ_4_ backbone dihedral angles, where one mode overlapped with the distribution of angles sampled in MD simulations of the 10LH-bound complex (Fig. 3b). These data suggest that the PHOX2B peptide can adopt two conformational states in the HLA-A*24:02 groove, where one of the states presents R6 in an optimal conformation for nanomolar-range binding to 10LH.

**Figure 3.**
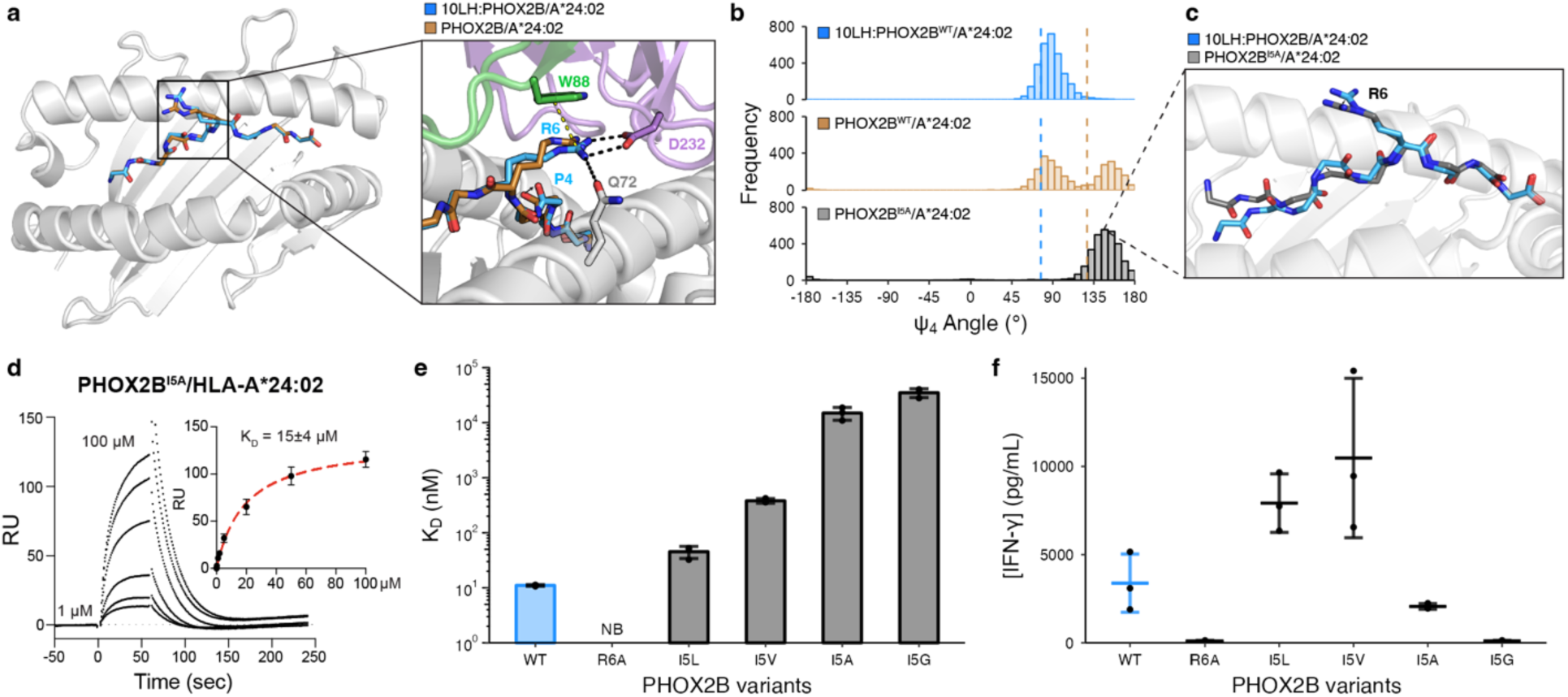
PHOX2B conformational adaptations enable high affinity complex formation with 10LH. **a,** Structural superimposition of PHOX2B/HLA-A*24:02 in the bound (blue; PDB ID 8EK5) and unbound (brown; PDB ID 7MJA) states. The peptides were aligned using their backbone heavy atoms (RMSD = 0.52 Å). The zoom-in details the difference in the R6 side chain conformation and the peptide backbone. **b,** Histogram depicting the frequency of Ψ_4_ backbone dihedral angle values for MD simulations of the bound (blue) and unbound (brown) PHOX2B/HLA-A*24:02 structures as well as the PHOX2B^I5A^/HLA-A*24:02 unbound variant (black). The dashed lines correspond to the values in the crystal structures (10LH:PHOX2B/HLA-A*24:02/β_2_m, PDB ID 8EK5; PHOX2B/HLA-A*24:02, PDB ID 7MJA). Data represent 3000 equally-spaced frames from a sum of n = 3 independent 1 μs runs. **c,** Representative structure from a frame of PHOX2B^I5A^/HLA-A*24:02 (black) MD simulations superimposed onto the 10LH:PHOX2B/HLA-A*24:02/β_2_m (blue) crystal structure. The peptides were aligned using all atoms (RMSD = 1.19 Å). **d**, Representative SPR sensorgram of PHOX2B^I5A^/HLA-A*24:02 flowed over a streptavidin chip coupled with 10LH-biotin. The concentrations of analyte for the top and the bottom sensorgrams are noted. Data are mean ± SD for n = 3 technical replicates. Fits from the kinetic analysis are shown with red lines. K_D_, equilibrium constant; RU, resonance units. **e,** SPR determined K_D_ values for HLA-A*24:02 loaded with I5 mutational scanned peptides. Data are mean ± SD for n = 3 technical replicates. NB, no binding. **f,** IFN-γ levels in supernatant of 10LH.BBz CAR T cells co-cultured with HLA-A*24:02 colorectal adenocarcinoma cell line SW620 pulsed with PHOX2B peptide variants. Exogenous peptide was added to SW620 target cells (15 µM final concentration). After four hours of incubation, 10LH.BBz CAR T cells were added to target cells at a 3:1 ET ratio. Supernatant was collected 24 hours post effector cell addition and IFN-γ levels were measured by ELISA. Values represent mean ± SD using effector cells from n = 3 biological donors, in triplicates.

The observed structural plasticity at position 4 of PHOX2B led to the hypothesis that presentation of the R6 sidechain could be influenced by the adjacent Ile residue at position 5, which forms hydrophobic contacts with the floor of the HLA-A*24:02 groove. MD simulations of the PHOX2B^I5A^/HLA-A*24:02 complex sampled a narrower distribution of Ψ_4_ angles relative to wild-type, limited to conformations that largely exclude those found in simulations of the 10LH-bound structure (Fig. 3b). Analysis of representative MD snapshots revealed a 1.4 Å shift of the P4 Cα atom, causing the peptide backbone to sink deeper into the HLA groove (Fig. 3c). This led to a 2.1 Å shift in the Cζ atom of R6, away from the optimal geometry for interactions with the 10LH CDRH3s. In agreement, the I5A substitution reduced binding to 10LH by approximately 1000-fold as determined by SPR (Fig. 3d, Extended Data Fig. 10, and Supplementary Fig. 6). These results suggest that peptide backbone structural adaptations, as sampled by the apo pHLA state, can drastically impact CAR-T recognition by affecting R6 rotamer selection and its priming for interactions with the scFv.

Next, we conducted MD simulations followed by SPR binding experiments, introducing Leu, Val, or Gly substitutions at position 5. We observed a negative correlation between the frequency of sampling Ψ_4_ angles, which resembled the scFv bound state (Extended Data Fig. 11a), and 10LH binding affinity, where conservative substitutions of I5 to Leu or Val maintained nanomolar-range K_D_ values (Fig. 3e, Extended Data Fig. 11b-e, and Supplementary Fig. 7). In contrast, I5A and I5G perturbation of R6 sidechain placement led to approximately 1,500- and 3,500-fold reduction, respectively, in 10LH binding affinity by sampling an altered peptide backbone conformation. Finally, we measured cytokine release triggered on 10LH CAR-T cells by peptide pulsing on HLA-A*24:02 positive antigen presenting cells (APCs) (Supplementary Fig. 8). Noticeably, although the SPR-determined K_D_ values of I5L and I5V were more than 4- and 30-fold lower than the WT, respectively, pulsed APCs with either mutant peptide demonstrated enhanced T cell activation relative to the WT peptide (Fig. 3f). This enhancement could be explained by either a difference in peptide loading or by our observed 10LH dissociation rate constants from the pHLAs, with k_d_ values of 0.0007 sec^-1^ and 0.0026 sec^-1^ versus 0.0022 sec^-1^ measured for I5L, I5V, and I5, respectively (Extended Data Table 3), in agreement with kinetic proofreading models for T cell receptor activation^25, 26^. In an analogous manner to TCRs, the contribution of both antigen density and pHLA dissociation rate for effective PHOX2B antigen discrimination by 10LH CAR-T cells may explain how HLA-A*23:01, which has approximately two orders of magnitude reduced binding to 10LH relative to HLA-A*24:02 (2.6 μM vs. 11 nM K_D_) shows potent CAR-T killing of tumor cells^2^, since both interactions show similar k_d_ values by SPR (0.0077 sec^-1^ versus 0.0022 sec^-1^). These results allow us to establish a relationship between pHLA dissociation rate from the scFv, known as CAR dwell time, and CAR-T cell activation, cytokine release, and cytotoxicity, established by previous studies focusing on both TCRs and CARs^27–29^.

### Molecular mimicry of the PHOX2B antigen elicits 10LH cross-reactivity with self-epitopes

To characterize the peptide cross-reactivity range of 10LH experimentally, we first applied a peptide X-scan^30^ approach to explore all amino acid (excluding Cys) substitutions of non-anchor PHOX2B residues and measured 10LH binding *in vitro* using a high-throughput assay, which leverages micro-refolding to generate fluorophore-labeled pMHC complexes^31^ (Fig. 4a). In agreement with our structural and SPR data (Extended Data Fig. 10), R6 is required while the I5L and I5V mutations can be tolerated for binding to 10LH, respectively (Extended Data Fig. 12a). We then generated a peptide pattern to capture the per-position amino acid binding preferences established from our X-scan assay, and obtained a set of 38 potentially cross-reactive peptide sequences found in the normal human proteome after cross-referencing with ScanProsite^32^ (Extended Data Fig. 12b). We repeated micro-refoldings with these peptides and confirmed binding to 10LH by SPR for three peptides in our set (ATG2A, CNGB3, and GLB1) in addition to two cross-reactive peptides identified previously by our selective cross-reactive antigen presentation (sCRAP) algorithm (ABCA8 and MYO7B)^2^. All peptides bind 10LH with K_D_ values ranging from 1 μM to 27 μM (Extended Data Fig. 13 and Supplementary Fig. 9), however, the ATG2A, CNGB3, and GLB1 peptides exhibit a slow dissociation rate, within the same profile as the cognate PHOX2B antigen. Focusing on the highest affinity peptide (ATG2A, LYLPVRVLI), we determined two complementary ATG2A/HLA-A*24:02 structures at 1.8 Å and 3.0 Å resolution (Extended Data Table 1). The 1.8 Å resolution structure showed one pHLA molecule in the asymmetric unit, where the key R6 side chain was buried in the HLA groove and inaccessible for 10LH recognition (Extended Data Fig. 14a). However, in the 3.0 Å resolution structure, one of the four pHLA complexes in the asymmetric unit adopted a peptide backbone which resembles PHOX2B with a solvent-exposed R6 side chain (Fig. 4c and Extended Data Fig. 14b-e). These results suggest that ATG2A/HLA-A*24:02 can sample a minor conformational state with a PHOX2B-like backbone structure as a potential basis for driving strong cross-reactivity with 10LH *in vitro*.

**Figure 4.**
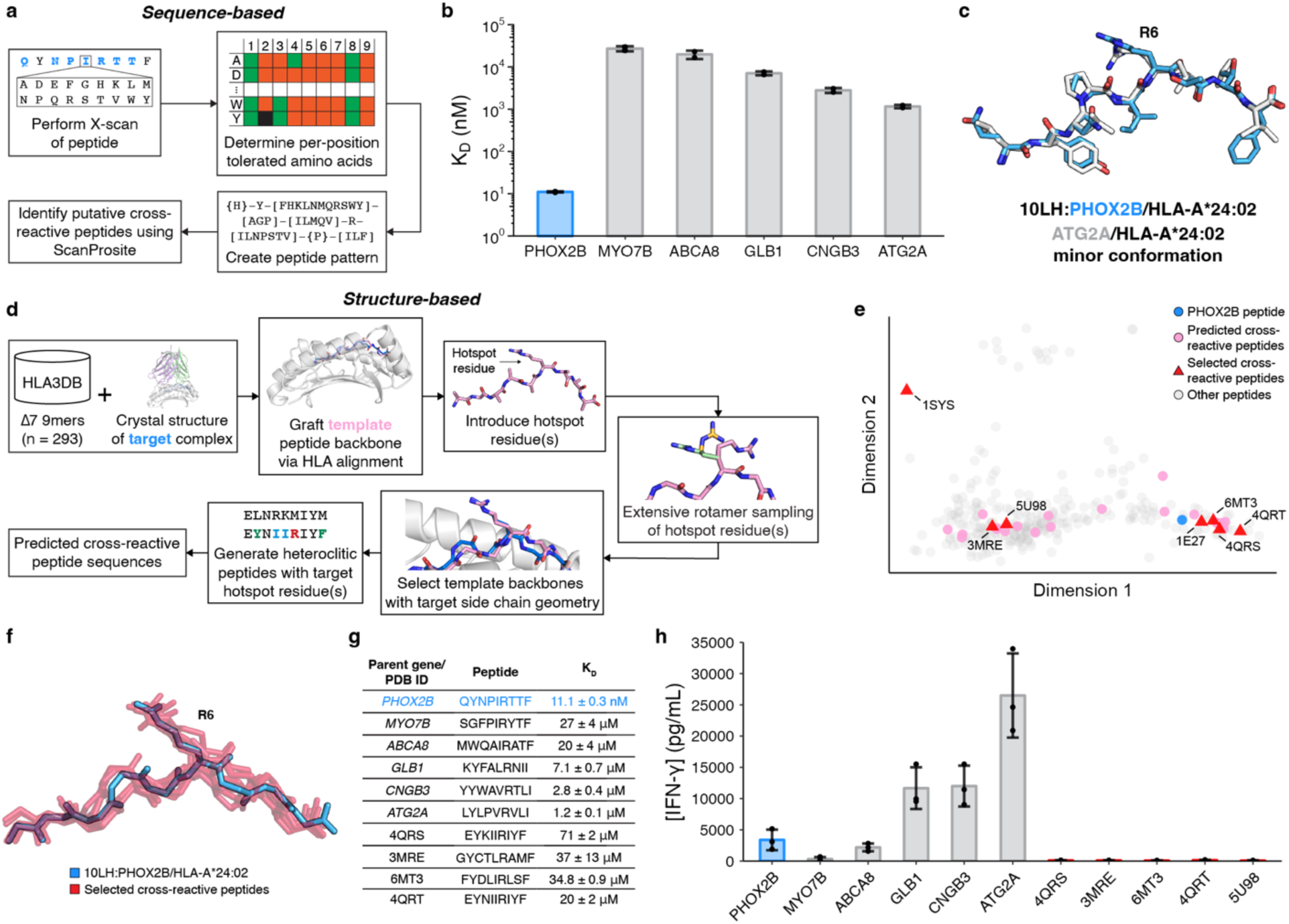
Mapping the 10LH/peptide cross-reactivity landscape. **a,** Summary of sequence-based prediction of cross-reactive peptides by X-scan/ScanProsite: (1) perform X-scan on target peptide; (2) determine per-position tolerated amino acids; (3) create peptide pattern; (4) identify putative cross-reactive peptide using ScanProsite. **b,** SPR determined K_D_ values for cross-reactive peptides. Data are mean ± SD for n = 2 technical replicates. **c**, Structural superimposition of the 10LH:PHOX2B/HLA-A*24:02/β_2_m peptide (blue; PDB ID 8EK5) with the minor conformation of the 3.0 Å structure of ATG2A/HLA-A*24:02 (white; PDB ID 8SBL). The peptides were aligned using their backbone heavy atoms (RMSD = 0.86 Å). **d,** Summary of structure-based prediction of cross-reactive peptides: (1) graft template peptide backbones from HLA3DB; (2) introduce hotspot residues; (3) conduct extensive rotamer sampling; (4) select template backbones with target side chain geometry; (5) generate heteroclitic peptides with target hotspot residues and conserved backbone; (6) obtain a set of predicted cross-reactive peptides. **e,** PCA plot of the conformational landscape of peptide backbones. The PCA transformation explains 66% of the variation. **f,** Structural superimposition of selected, predicted cross-reactive peptide backbones (red) to the PHOX2B backbone (blue) with the side chain of R6 shown. The peptides were aligned using HLA Cα atoms. **g,** Table of cross-reactive peptide sequences with their origin and SPR-determined K_D_ values. Data are mean ± SD for n = 2 or 3 technical replicates. **h,** IFN-γ levels in supernatant of 10LH.BBz CAR T cell co-cultured with HLA-A*24:02 colorectal adenocarcinoma cell line SW620 pulsed with cross-reactive peptides. Exogenous peptide was added to SW620 target cells (15 µM final concentration). After four hours of incubation, 10LH.BBz CAR T cells were added to target cells at a 3:1 ET ratio. Supernatant was collected 24 hours post effector cell addition and IFN-γ levels were measured by ELISA. Values represent mean ± SD using effector cells from n = 3 biological donors, in triplicate.

To further explore the conformational landscape of possible cross-reacting peptides from a structural perspective, we leveraged our annotated database of high-resolution pHLA structures, HLA3DB^33^ to search for complementarity with the 10LH recognition mode. We used our high-resolution structure of the 10LH:PHOX2B/HLA-A*24:02/β_2_m complex as a modeling template, aligned the HLA Cα atoms from all existing pHLA complexes to the scFv-bound state of PHOX2B, and performed extensive rotamer sampling of an introduced R6 residue, aiming to identify backbone configurations which allowed optimal side chain placement for interactions with 10LH CDRs (Fig. 4d). We identified 29 potentially cross-reactive peptide backbone scaffolds spanning the observed conformational landscape (Fig. 4e, f). We selected 7 representative peptides, and further optimized for binding to HLA-A*24:02 and 10LH through the introduction of anchor residue substitutions and the I5/R6 recognition motif, respectively. SPR experiments showed that 10LH could bind to 6 peptides in our set with K_D_ values in the range of 20 μM to 200 μM, yet exhibited a moderate to rapid dissociation rate (Fig. 4g, Extended Data Fig. 15, Supplementary Fig. 9, and Extended Data Table 3). These data indicate that 10LH can engage antigens with a non-PHOX2B-like backbone, albeit with a significant reduction in binding affinity.

To evaluate the functional consequences of the *in vitro* identified peptide cross-reactivity, we measured cytokine release upon CAR-T stimulation with soluble peptides pulsed on HLA-A*24:02 positive APCs. We observed enhanced peptide-dependent cytokine release relative to the WT peptide for three out of five antigens obtained from our sequence-based approaches(Fig. 4h and Supplementary Fig. 10). Consistently, this can be rationalized by an improvement in the dissociation rate (Extended Data Table 3), in accordance with kinetic proofreading models for T cell receptor activation^25–29^. The differences observed were not a result of pMHC complex stability (Supplementary Fig. 9). While peptide cross-reactivity was observed in this cytokine release assay, our prior study^2^ indicated that these antigens (ATG2A, CNGB3, and GLB1) are not of clinical concern; cell lines expressing the putative cross-reactive proteins showed no cytotoxicity or IFN-γ release, suggesting that the cross-reactive peptides were not presented on the cell surface. Meanwhile, antigens derived from our orthogonal, structure-guided approach had a moderate to fast dissociation rate and thus did not present a cross-reactivity risk as shown by a lack of sufficient cytokine release (Extended Data Table 3). These data suggest that while all *in vitro* identified cross-reactive peptides can form an initial encounter complex^34^ with 10LH, guided by the R6 electrostatic interaction and contacts with HLA framework residues, only a subset can transition to an ideal conformation that produces downstream signaling as measured by IFN-γ release. One possible mechanism for this transition is through intrinsic peptide dynamics, enabling R6 structural adaptations to drive high-affinity complex formation and ultimately leading to slow dissociation rate constant which triggers signaling. This possible mechanism can provide an additional fidelity checkpoint which limits off-target CAR-T cell activation by self-antigens. Taken together, these findings establish that 10LH triggering on CAR-T cells necessitates not only a specific peptide backbone conformation, thereby achieving sufficient CAR engagement, but also a slow dissociation rate achieved through peptide dynamics, which leads to a narrow cross-reactivity profile for 10LH. Moreover, the only strong cross-reactivity identified from our comprehensive search did not correspond to a risk as the putative peptide was not presented in healthy tissue.

## Discussion

Through a combination of structural, biochemical, and functional approaches, we have characterized the molecular basis for peptide-centric recognition of HLA molecules by the 10LH PC-CAR. Our work delineates important structural principles. We find that the formation of a high-affinity complex necessitates a set of conserved HLA framework residues, restricting clinical application to patients expressing an HLA allotype from the A9 serological CREG. Additionally, conformational plasticity of the peptide highlights the importance of sampling a specific backbone structure for CAR-T triggering and cytokine release. While we identify three putative cross-reactive peptides, their lack of expression in cell lines indicates that they do not pose a real-world risk. Taken together, our analysis reveals significant advantages of PC-CARs over traditional T-cell receptor immunotherapies. As TCRs are inherently cross-reactive^35, 36^ their micromolar-range affinity allows them to recognize different epitopes through a variety of strategies^37^ not limited to differential docking^38^ and accepting structural adjustments^39^. In contrast, by adopting a restricted, converged binding geometry with a specific peptide backbone for high-affinity complex formation, 10LH allows for a more restricted antigen cross-reactivity profile, which mitigates off-target toxicity liabilities relative to conventional TCRs.

In contrast to TCR engineering approaches^40^, PC-CAR development can yield nanomolar-range, peptide-specific scFv binding without the need for further affinity enhancement. Thus, our binding mechanism is well-poised for future development of bispecific T cell engager constructs (BiTEs)^41, 42^. In addition, our results provide a roadmap for repurposing^43^ 10LH to target the PHOX2B antigen displayed by other common HLA alleles that are not part of the A9 CREG, such as HLA-B*14:02. Alternatively, as shown recently for other MHC-I antibodies^43^, our structure provides a basis for expanding the scope of peptide targets presented by HLA-A*24:02 through the rational design of the CDR3 loops. Thus, the structural principles established through our study can be leveraged to develop a library of nanomolar-range binders that address multiple tumor-associated peptides and HLA specificities, providing optimal modalities for personalized cancer immunotherapy.

### Methods Crystallography

The 10LH:PHOX2B/HLA-A*24:02/β_2_m complex was prepared by mixing PHOX2B/HLA-A*24:02/β_2_m and 10LH at 1.5:1 molar ratio and incubated for 1 hour at 4 °C. Complex was purified by size exclusion chromatography (Supplementary Fig. 1a). The 10LH:PHOX2B/HLA-A*24:02/β_2_m complex at 9.5 mg/ml was used for growing crystals and crystals were obtained in 0.2 M Sodium nitrate and 20% w/v PEG 3350 at 1:1 ratio. Data was collected at SSRL beamline BL 12-2 using Dectris Pilatus 6M detector (wavelength, 0.979 Å) and diffraction data was processed using XDS (v. Jan 10, 2022)^44^. Structure of the 10LH:PHOX2B/HLA-A*24:02/β_2_m complex was solved by molecular replacement method using Phenix (v. 1.20.1-4487)^45^ and Refmac5 software (v. 5.5)^46^ of CCP4 (v. 8.0) package suites^47^ for refinement and COOT (v. 0.9.8.4)^48^ was used for model building. PDB-REDO was used for further model refinement^49^. Favored/allowed/outlier for the structures are 98/2/0. The CDR loops were defined using the Chothia definition^50^ as implemented in abYsis^51^.

For ATG2A/HLA-A*24:02, a sample at 14.4 mg/ml was mixed with reservoir well solution at 1:1 ratio and incubated at 20 °C. Crystals were obtained in two conditions, 50 mM MES monohydrate pH 6.5 and 22.5% w/v PEG 3350, and 2 M Ammonium sulfate and 0.1 M Sodium cacodylate pH 6.5, which diffracted at 3 Å and 1.8 Å, respectively. Diffraction data were collected at BNL, NSLS II, 17-ID-1 (AMX) using Dectris Eiger 9M detector at a wavelength of 0.92 Å. XDS^44^ software was used for data processing and structures were solved using Phaser (v. 2.8.3)^52^ and Buster Global Phasing^53^ or Phenix^45^. Ramachandran statistics for the structures showed the percentage of residues in favored/allowed/outliers were, 99/1/0 and 90/8/2, respectively.

### AlphaFold structure prediction

The AlphaFold models of the 10LH:PHOX2B/HLA-A*24:02 complex were calculated with AlphaFold^16^ (v. 2.3.1) using the multimer model configuration in the non-docker setup (https://github.com/kalininalab/alphafold_non_docker)^16,54^. The full genetic database configuration was used for multiple sequence alignments and post-prediction relaxation of all models was achieved using the default method, gradient descent in the Amber force fields^55^. Models were generated using a maximum template release date of 2022-01-06.

### Comparison to TCR structures

The TCR:nonamer/HLA-I structural dataset was generated using a modified version of HLA3DB^33^, which identifies structures in the Protein Data Bank (PDB)^56^ using the RCSB PDB Search API (v. 2)^57^. Using the same selection criteria as HLA3DB, we obtain a total of 67 crystal structures (as of June 9^th^, 2022). Complexes were analyzed using PDBePISA^23^ as implemented in CCP4 (v. 8.0)^47^ to obtain peptide/receptor and HLA/receptor interface area values and interaction residues. The corresponding TCR docking angles were obtained or calculated from TCR3d^58^.

### Molecular dynamics simulations

All-atom molecular dynamics (MD) simulations in explicit solvent were carried out as previously described in GROMACS (v. 2020.4)^59^ using an AMBER99SB-ILDN protein forcefield and TIP3P water model^60^. LINCS and SETTLE constraint algorithms were used to constrain peptide/protein and water molecules, respectively. An integration time step of 2 fs was used with coordinates output every 10 ps. Short-range interactions were treated with a Verlet cut-off scheme with 10 Å electrostatic and van der Walls cutoffs and long-range electrostatics were treated with the PME method with a grid spacing of 1.2 Å and cubic interpolation. Simulations were carried out in cubic simulation boxes, and periodic boundary conditions were used in all three spatial dimensions. The thermodynamic ensemble was NPT with temperature was kept constant at 300 K by a velocity rescaling thermostat with a stochastic term^61^ with 0.1 ps time constant and pressure kept constant at 1 bar pressure using an isotropic Berendsen barostat with 0.5 ps time constant and 4.5 × 10^−5^ bar^−1^ isothermal compressibility. The 10LH:PHOX2B/HLA-A*24:02/β_2_m crystal structure (PDB ID 8EK5) served as the primary input with the HLA-A*24:02 α_3_ domain and β_2_m portions removed to ease compute time. For modelling of PHOX2B peptide mutations, Maestro (Schrödinger) was used to generate mutations in the peptide sequence. Each system was solvated to overall neutral charge and contained Na^+^ and Cl^−^ ions to yield physiological concentration of 0.15 M. Following 500 steps of steepest-descent energy minimization, initial velocities were generated at 65 K with linear heating up to 300 K over 2 ns. Three independent trajectories for each scFv:pHLA-I and pHLA-I complex were acquired for 1 μsec using the Children’s Hospital of Philadelphia Respublica computational cluster. Analysis was performed using GROMACS and the Visual Molecular Dynamics (VMD) package^62^. Visualizations were produced with PyMOL (v. 2.5.2)^63^, R (v. 4.2.1), and RStudio (v. 2023.03.0+386).

For analysis, periodic boundary conditions were removed and 1000 equally spaced frames from each trajectory (3000 frames in total for three independent replicates) were extracted in using `gmx trjconv`. Phi/psi/chi angles and dihedral order parameters were calculated using `gmx chi`.

Peptide per-atom RMSF was calculated for each individual replicate and averaged across replicates using `gmx rmsf`. A frame from the PHOX2B^I5A^/HLA-A*24:02 was extracted by selecting a representative frame within a D-score^33^ of 0.1 from the median value of the ψ_4_ angle for this distribution (150°) using a custom Python script.

### Peptides

A full list of the HLA peptides used in this study is shown in Supplementary Table 1. All peptide sequences were given as standard single-letter codes and were purchased from Genscript, NJ, USA, at >90% purity. For the peptide solutions, lyophilized peptides were solubilized in distilled water and centrifuged at 14,000 rpm for 15 min. Concentrations were calculated using the respective absorbance and extinction coefficient at 205 nm wavelength. X-scan peptides were purchased from Mimotopes (Victoria, Australia).

### Recombinant protein expression, refolding, and purification

Plasmid DNA encoding the BirA substrate peptide (BSP, LHHILDAQKMVWNHR)-tagged luminal domain of HLA heavy chains and human β_2_m were provided by the NIH tetramer facility (Emory University) and additional HLA encoding constructs were cloned into pET-22b(+) vector using NdeI/BamHI restriction sites (Genscript) or produced through site-directed mutagenesis. Plasmids were transformed into *Escherichia coli* BL21(DE3) cells (New England Biolabs), expressed in Luria Broth and inclusion bodies were solubilized using guanidine hydrochloride as previously described^64^. pHLA-I complexes were generated by *in vitro* refolding as 200 mg mixtures of heavy chain:light chain at a 1:3 molar ratio and 10 mg of peptide in 1 L of refolding buffer (0.4 M L-Arginine-HCl, 2 mM EDTA, 4.9 mM reduced L-Glutathione, 0.57 mM oxidized L-Glutathione, 100 mM Tris pH 8.0) at 4 °C. Refolding proceeded for 4 days and the pHLA-I complex was purified by size-exclusion chromatography (SEC) using a HiLoad 16/600 Superdex 75 pg column at 1 mL/min with 150 mM NaCl, 25 mM Tris buffer, pH 8.0. The luminal domain of the TCR NYE-S1 α/β complex was expressed and purified as previously described^65^. The 10LH scFv protein was provided by Myrio Therapeutics (Australia). Protein concentration was determined using A_280_ measurements on a Nanodrop with extinction coefficients estimated with ExPASy ProtParam tool^66^.

### Differential Scanning Fluorimetry

Samples were prepared at a final concentration of 7 μM in PBS buffer (150 mM NaCl, 20 mM sodium phosphate pH 7.4) and mixed with 10X SYPRO Orange dye (ThermoFisher) to a final volume of 20 μL. Samples were then loaded into a MicroAmp Fast 384-well plate and ran in triplicates (n = 3) on a QuantStudio™ 5 Real-Time PCR machine with excitation and emission wavelengths set to 470 nm and 569 nm, respectively. Temperature was incrementally increased at a rate of 1 °C/min between 25 °C and 95 °C to measure the thermal stability of the proteins. Data analysis and fitting was performed in GraphPad Prism v9.

### Biotinylation and Tetramer formation

Biotinylation and tetramer formation of the 10LH molecules were performed as previously described^67^. In brief, BSP-tagged 10LH proteins were biotinylated using the BirA biotin-ligase bulk reaction kit (Avidity), according to the manufacturer’s instructions. Streptavidin-PE (Agilent Technologies, Inc.) at 4:1 monomer: streptavidin molar ratio was added to the biotinylated pHLA-I in the dark, every 10 min at room temperature over 10-time intervals.

### Surface plasmon resonance

SPR experiments were conducted in duplicates or triplicates (n = 2 or 3) using a BiaCore X100 instrument (Cytiva) in SPR buffer (150 mM NaCl, 20 mM sodium phosphate pH 7.2, 0.1% Tween-20). Approximately 2000 resonance units (RU) of biotinylated-10LH were immobilized at 10 µL/min on a streptavidin-coated chip (Cytiva). Samples of pHLA-I at various concentrations were injected over the 10LH-streptavidin-coated chip at 25 °C at a flow rate of 30 µL/min for 60 sec followed by a buffer wash with 180 sec dissociation time. Equilibrium data were collected. The SPR sensorgrams, association/dissociation rate constants (k_a_, k_d_), and equilibrium dissociation constant K_D_ values were analyzed in BiaCore X100 evaluation software (Cytiva) using kinetic analysis settings of 1:1 binding or fitted using one-site specific binding (affinity fit) by GraphPad Prism v9. SPR sensorgrams and affinity-fitted curves were prepared in GraphPad Prism v9.

### HLA sequence analysis

Global HLA allele frequencies were obtained from the Allele Frequency Net Database^68^ and converted to population frequencies assuming a Hardy-Weinberg equilibrium and ignoring linkage. For alleles with a population frequency greater than 0.1% (“common allele,” n = 346), sequences were obtained from the IMGT/HLA database^69^ and a pairwise sequence alignment was performed to the HLA-A*02:01 heavy chain sequence (consisting of 180 residues from the

N-terminus). Then, NetMHCPan 4.1^20^ was utilized to determine strong (≤ 0.5% rank) and weak (≤ 2% rank) PHOX2B-binding alleles (n = 219) for further analysis. American allele frequencies were obtained from the BeTheMatch registry^70^ across five broad race groups and converted to population frequencies assuming a Hardy-Weinberg equilibrium and ignoring linkage. A combined American population frequency for each allele was obtained by normalizing values according to the racial distribution followed by the 2020 U.S. Census. Alleles with a total population frequency greater than 0.01% were considered.

### SPR binding penalty

For a given allele, an SPR binding penalty was determined by adding the log base 10 ratio of the mean K_D_ value for each polymorphism to the wild-type K_D_ (11 nM). For polymorphisms that resulted in no observable binding (e.g. G65R), an artificial K_D_ of 1200 µM was utilized which added 5 to the SPR binding penalty. Calculations were implemented using a custom Python script.

### Structural modeling

To model different alleles onto the 10LH:PHOX2B/HLA-A*24:02/β_2_m structure, we utilized the RosettaRemodel^71^ application from the Rosetta (v. 2020.08)^72^ suite of programs. All side chains of the template structure aside from those modeled on the HLA were left in their original poses. A blueprint file was utilized to specify the polymorphisms relative to the HLA-A*24:02 sequence.

### Single-antigen bead assay

Single antigen beads (SABs) are fluorescently color-coded and coated with 97 different HLA allotypes^73, 74^, derived from Epstein-Barr virus (EBV)-transfected cell lines and are, thus, loaded with a pool of different cell-derived peptides. The levels of pHLAs across the beads were consistent, as confirmed by staining with the pan-HLA class I W6/32 antibody (Extended Data Fig. 8a). Loading of either PHOX2B or a negative control peptide (PHOX2B^R6A^) on a subset of the HLAs with the corresponding peptide-binding specificities is mediated under suitable conditions (Extended Data Fig. 8b). Bead binding experiments employ tetramerized 10LH, coupled to an orthogonal fluorophore, for probing interactions with HLAs with a high-to-intermediate affinity, up to micromolar K_D_ range (Fig. 2h). To test the levels of peptide-loaded HLA-I molecules on the SABs, we used a PE-conjugated W6/32 antibody (Biolegend, 311406) mixed with 4 μL of the LABScreen single antigen HLA-I bead suspension (OneLambda, Inc., CA, USA) in a 96-well plate. The samples were incubated for 30 min, 550 rpm at room temperature (RT), washed four times in Wash Buffer (OneLambda, Inc., CA, USA) to remove excess of antibody and resuspended in phosphate-buffered saline (PBS), pH 7.2. To screen for low-affinity interactions, we generated 10LH tetramers by conjugating biotinylated 10LH monomer with Streptavidin-PE (Agilent Technologies, Inc.) at 4:1 monomer/streptavidin molar ratio as previously described^75^. 80 nM of 10LH in tetramers were mixed with 4 μL of beads pre-incubated o/w with 100-fold molar excess of the desired peptides, incubated for 1h, 550 rpm at RT followed by four washes and resuspension in PBS buffer. In all cases, the levels of fluorescence intensity were measured using the Luminex 100 Liquid Array Analyzer System and the results were analyzed in GraphPad Prism v9.

### X-Scan/ScanProsite bead binding assay

For each non-anchor position of the PHOX2B epitope, peptides were synthesized (Mimotopes, Australia) representing all amino acids except cysteine. Each peptide was micro-refolded with BFP-ligated MHC alpha-chain and fluorophore-labeled β_2_m to form fluorophore-labeled pMHC complexes^31^. Streptavidin C1 Dynabeads (Life Technologies) were incubated with excess biotinylated scFv before being blocked with free biotin and washed in SMG (100 mM NaCl, 8 mM MgSO_4_ 7H_2_O, 50 mM Tris-HCl, pH 7.5). Each fluorophore-labeled pMHC complex was added to a concentration of 3.5 nM and incubated for one hour at 4 °C followed by ten minutes at 25 °C. Binding of the free MHC complex to the beads was quantified by CytoFLEX (Beckman Coulter) at 488 nm (excitation)/525 nm (emission). Binding was normalized to beads without scFv and with unrelated control MHC complex. An amino acid was deemed to be tolerated if it had 15% of the fluorescence of the PHOX2B peptide. Tolerated residues were used to generate the peptide pattern, {H}-Y-[FHKLNMQRSWY]-[AGP]-[ILMQV]-R-[ILNPSTV]-{P}-[ILF], which was used to query ScanProsite. The one-letter code in square brackets “[ ]” indicates acceptable amino acids and in curly brackets “{ }” indicates amino acids not accepted at that position. Potential off-target peptides identified by the ScanProsite query were synthesized (Mimotopes) micro-refolded and tested for 10LH scFv binding under the same conditions as the X-scan. The top three peptides, ranked by mean fluorescence relative to the PHOX2B peptide, were selected for further analysis.

### Cell lines

Human neuroblastoma cell line SK-N-DZ was obtained from the Children’s Hospital of Philadelphia Maris cell line bank and was cultured in RPMI supplemented with 10% fetal bovine serum (FBS), 100 U/mL penicillin, 100 µg/mL streptomycin, and 2 mM L-glutamine. Colorectal adenocarcinoma cell line SW620 was obtained from American Type Culture Collection (ATCC) and was cultured in RPMI supplemented with 10% FBS, 100 U/mL penicillin, 100 µg/mL streptomycin, and 2 mM L-glutamine. Packaging cell line HEK 293T was obtained from ATCC and was cultured in DMEM supplemented with 10% FBS, 100 U/mL penicillin, 100 µg/mL streptomycin, and 2 mM L-glutamine. All cell lines were grown under humified conditions in 5% CO_2_ at 37 °C, and samples were regularly tested for mycoplasma contamination.

High resolution class I HLA typing was performed by the Children’s Hospital of Philadelphia Immunogenetics Laboratory.

### Primary human T cells

Primary human total T cells (CD4^+^/CD8^+^) were obtained from anonymous donors through the Human Immunology Core at the Perelman School of Medicine at the University of Pennsylvania under a protocol approved by the Children’s Hospital of Philadelphia Institutional Review Board. Cells were cultured using AIM-V (Thermo Fisher Scientific) supplemented with 10% FBS, 100 U/mL penicillin, 100 µg/mL streptomycin, and 2 mM L-glutamine under humified conditions in 5% CO_2_ at 37 °C.

Primary human T cells were thawed and activated in culture for 1 day in the presence of 5 ng/ml recombinant IL-7, 5 ng/ml recombinant IL-15, and anti-CD3/CD28 beads (Dynabeads, Human T-Activator CD3/CD28, Life Technologies) at a 3:1 bead:T cell ratio in G-Rex system vessels (Wilson Wolf). On day 2, thawed lentiviral vector was added to cultured T cells with 10 µg/mL Polybrene (Millipore Sigma), and 24 hours later, vessels were filled with complete AIM-V medium supplemented with indicated concentrations of IL-7 and IL-15. On day 10 cells were harvested, washed, and activation beads were magnetically removed. Cells were cultured until day 12, at which transduction efficiency was determined using flow cytometry and cells were frozen (Supplementary Fig. 5).

### Peptide pulsing assay

Target cells were plated in triplicate at 20E3 cells/well in 96-well tissue culture treated plates. CAR T cells and mock T cells were thawed and cultured overnight in 5 ng/ml IL-7 and 5 ng/ml IL-15. The following morning, exogenous peptide was added to target cells at a final concentration of 15 µM. Peptide concentration was confirmed prior to addition using a NanoDrop One spectrophotometer. Four hours later, washed CAR T cells and mock T cells were added at indicated E:T ratios to target cells. Supernatant was collected 24 hours post effector cell addition.

### xCELLigence RTCA cytotoxicity assay

Target cells were plated at 20E3 cells/well in triplicate or quadruplicate in 96-well Real-Time Cell Analysis (RTCA) E-Plates (Agilent). CAR T cells and mock T cells were thawed and cultured overnight in 5 ng/ml IL-7 and 5 ng/ml IL-15. The following day, CAR T cells and mock T cells were added at indicated E:T ratios to target cells. Target cell viability was determined using the cell index of the xCELLigence RTCA system. Target cell index was normalized to the timepoint immediately before addition of effector cells.

### Cytokine secretion assay

IFN-γ levels were determined from supernatant collected from peptide pulsing and cell cytotoxicity assays using ELISA kits according to the manufacturer’s protocol (BioLegend).

### Lentiviral generation

A second-generation lentiviral system was used to produce replication-deficient lentivirus. The day preceding transfection, 15 million HEK 293T cells were plated in a 15-cm dish. On the day of transfection, 80 µL Lipofectamine 3000 (Life Technologies, Invitrogen) was added to 3.5 mL room-temperature Opti-MEM medium (Gibco). Concurrently, 80 µL P3000 reagent (Thermo Fisher Scientific), 12 µg psPAX2 (Gag/Pol), 6.5 µg pMD2.G (VSV-G envelope), and a matching molar quantity of transfer plasmid were added to 3.5 mL room-temperature Opti-MEM medium. After five minutes of room-temperature incubation, the lipofectamine mixture was added dropwise to the plasmid mixture and incubated for 20 minutes at room temperature. Medium on the 293T plates was replaced with 13 mL of room-temperature Opti-MEM, after which the 7 mL transfection mixture was added dropwise. After 16-20 hours, the transfection medium was replaced with complete AIM-V medium. Virus supernatant was collected at 24 and 48 hours later, briefly centrifuged at 300 g, and passed through a 0.45 µM syringe. Supernatant was concentrated to 50–100X using Amicon 100 KDa centrifugal filter units (Millipore).

### Structure-based cross-reactive peptide prediction

The 10LH:PHOX2B/HLA-A*24:02/β_2_m crystal structure was utilized as the “target” complex with the β_2_m light chain removed and the first 180 residues of the HLA heavy chain utilized. Template pHLAs obtained from HLA3DB^33^ with a cutoff date of April 29^th^, 2022 were grafted onto the target complex via alignment of the HLA Cα atoms in PyMOL (v. 2.5.3)^63^. Then, the *fixbb* package in Rosetta (v. 2020.08)^47, 72^ was utilized to mutate the peptide sequence to alanine except for the R6 hotspot residue (AAAAARAAA) and conduct extensive χ angle rotamer sampling of the side chain peptide residues using the “-ex1 -ex2 -ex3 -ex4” parameters. To determine template backbones with ideal target side chain geometry, the distance between the R6 CZ atom and D232-CDRH3 CG atom was computed in PyMOL. The NH1-OD2 dihedral angle between the planes defined by (R6-CZ, R6-NH1, D232-OD2) and (R6-NH1, D232-OD2, D232-CG) was computed as well as the NH2-OD1 dihedral angle between the planes defined by (R6-CZ, R6-NH2, D232-OD1) and (R6-NH2, D232-OD1, D232-CG). Four criteria were used to determine if the side chain geometry was proper: (1) CZ-CG distance less than 5 Å; (2) D-score(10LH NH2-OD1, template NH2-OD1) less than 2.3; (3) D-score(10LH NH1-OD2, template NH1-OD2) less than 2.3; (4) Sum of D-score values less than 4.0. There were 29 template backbones that met this criterion, and the visual analysis confirmed that the side chain geometries were ideal. The sequence of the template backbone was altered to match the PHOX2B peptide characteristics. Specifically, (1) The anchor residues position 2 and position 9 are Tyr and Phe, respectively; (2) position 6 is Arg; (3) If position 4 or position 5 have positively charged side chains (R, H, or K), then the position is converted to Ile; (4) If position 7 has a positively charged side chains, it is converted to Gly. Thus, we obtained 25 peptide sequences as four template peptides converged upon alteration. A two-dimensional PCA plot was constructed by standardizing the peptide dihedral angles of position 4 to position 7 with the sine of the angle. A custom Python script was used for all analyses.

### Data Availability

Protein structures are available in the Protein Data Bank under accession codes 8EK5 (10LH:PHOX2B/HLA-A*24:02/β_2_m, QYNPIRTTF), 8SBK (ATG2A/HLA-A*24:02, LYLPVRVLI), and 8SBL (ATG2A/HLA-A*24:02, LYLPVRVLI). All other data are available within the article and supplementary information files, or by request from the corresponding author.

### Code Availability

Code used for the structure-based prediction of cross-reactive peptides can be accessed on GitHub via https://github.com/titaniumsg/pc_car. All other code is available upon reasonable request to the corresponding author.

## Acknowledgments

This work was supported in part by a St. Baldrick’s Foundation-Stand Up to Cancer Dream Team Translational Research Grant (SU2C-AACR-DT-27-17). The St. Baldrick’s Foundation collaborates with Stand Up To Cancer. Research Grants are administered by the American Association for Cancer Research, the Scientific Partner of SU2C. Stand Up To Cancer is a program of the Entertainment Industry Foundation administered by the American Association for Cancer Research (J.M.M.). This work was also supported by NIH grants U54 CA232568 as part of the Beau Biden Cancer Moonshot Program (J.M.M.) and NIH R35 CA220500 (J.M.M.), R01 AI143997 (N.G.S.), R35 GM125034 (N.G.S.), the Science Center Quod Erat Demonstrandum (QED) program at the Philadelphia Science Center (J.M.M. and M.Y.), the Children’s Hospital of Philadelphia Cell and Gene Therapy Collaborative (J.M.M. and M.Y.), the Giulio D’Angio Endowed Chair (J.M.M.), and Next Generation T Cell Therapies for Childhood Cancers – Cancer Grand Challenges (J.M.M. and N.G.S.).

This research used resources [AMX, 17-ID-1] of the National Synchrotron Light Source II, a U.S. Department of Energy (DOE) Office of Science User Facility operated for the DOE Office of Science by Brookhaven National Laboratory (BNL) under Contract No. DE-SC0012704. The Center for BioMolecular Structure is supported by NIH, National Institute of General Medical Sciences (NIGMS) through a Center Core P30 Grant (P30GM133893), and by the DOE Office of Biological and Environmental Research (KP1607011). This research also used resource of Stanford Synchrotron Radiation Lightsource (SSRL), SLAC National Accelerator Laboratory supported by the DOE, Contract No. DE-AC02-76SF00515. The SSRL Structural Molecular Biology Program is supported by NIH NIGMS (P30GM133894). We are grateful to Dr. Gino Cingolani for assistance with crystallographic refinement of the 10LH:PHOX2B/HLA-A*24:02/β2m complex.

## Author Contributions

Y.S., S.G., Q.F.M., J.M.M., and N.G.S. designed experiments. T.J.F. and L.M. purified 10LH complexes by SEC, performed all crystallization screening, diffraction data collection, model building and structure refinement. Y.S. performed all SPR experiments and data analysis. S.G. performed molecular dynamics simulations, structural modeling, 10LH vs. TCR structural analysis, and structure-based prediction of cross-reactive peptides. M.C.Y. designed plasmids, prepared all recombinant protein samples used in this study, and performed all DSF and quality control experiments. S.E.G. performed molecular dynamics simulations, data analysis and structure refinement of the final 10LH complex. G.F.P. developed the SAB cross-reactivity assay and performed all experiments and data analysis. H.V.T. performed preliminary protein crystallization screening. Q.F.M., E.M., and P.L. performed all CAR-T experiments and data analysis. A.F. and M.Y. obtained AlphaFold model of the 10LH complex. M.D.B., B.R.K., N.L.C., and S.J. prepared recombinant 10LH samples, performed peptide X-scan experiments and data analysis. S.G., Y.S., and N.G.S. wrote the paper, with feedback from all authors. J.M.M. and N.G.S. acquired funding and supervised the project.

## Corresponding author

Nikolaos G. Sgourakis;

## Ethics declarations

G.F.P., M.Y., Q.F.M., B.R.K., M.D.B., J.M.M., and N.G.S are listed as co-inventors in provisional patent applications related to this work. J.M.M., M.Y., Q.F.M., N.L.C., S.J., M.D.B., and B.R.K. have interests in companies involved in the commercialization of 10LH.

## Extended Data Figures

**Extended Data Figure 1.**
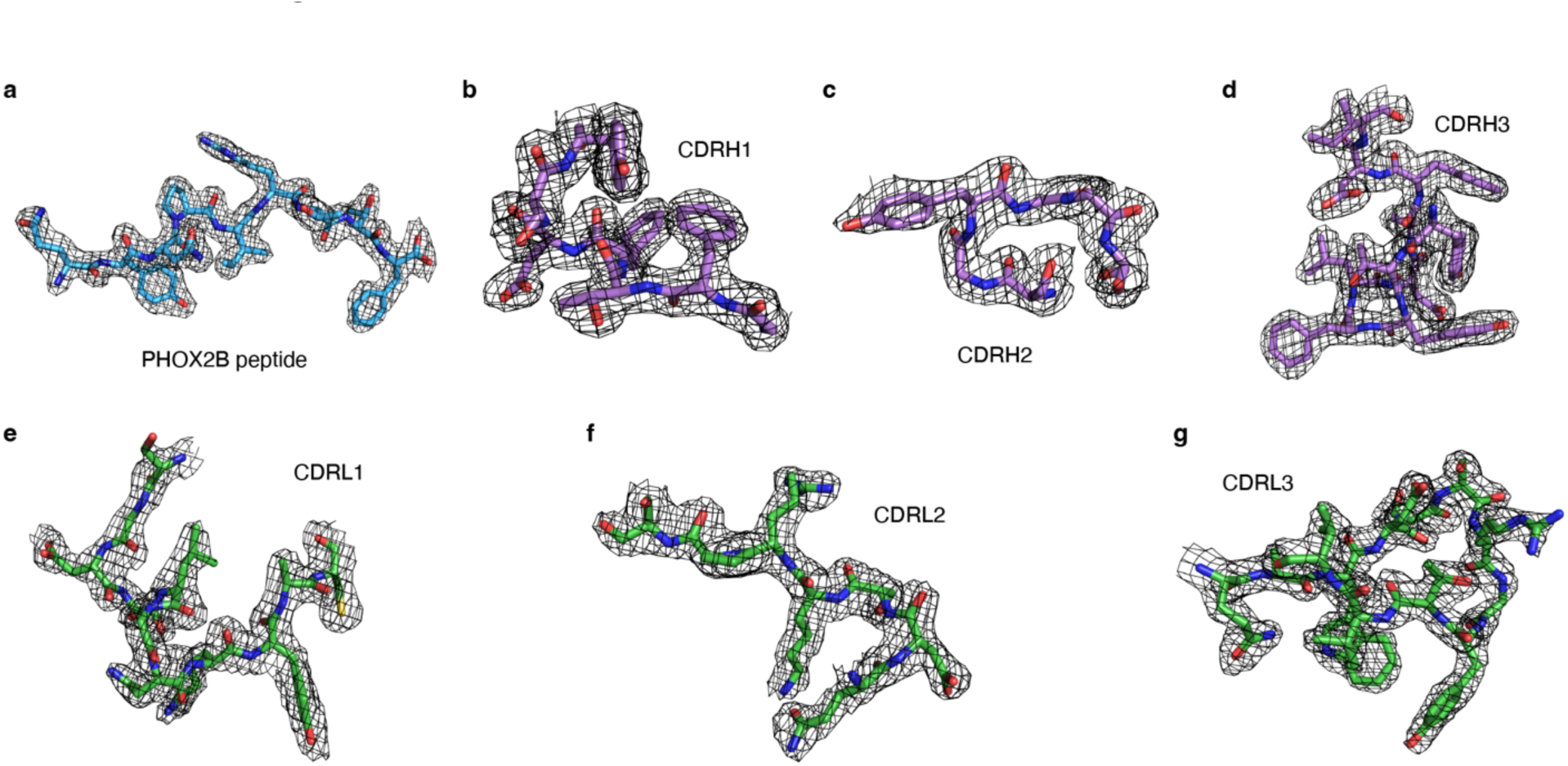
Structure determination of the 10LH:PHOX2B/HLA-A*24:02/β_2_m complex. **a**, Composite omit map of the PHOX2B peptide in the 10LH:PHOX2B/HLA-A*24:02/β_2_m complex (PDB ID 8EK5) with the backbone and side chains shown in blue sticks. The map is contoured at 1.1σ. **b-d,** Composite omit map of the CDRH1 (**b**), CDRH2 (**c**), and CDRH3 (**d**) of 10LH. The CDRH loops are shown in purple sticks. The map is contoured at 1.0σ for CDRH1, CDRH2, and CDRH3. **e-g,** Composite omit map of the CDRL1 (**e**), CDRL2 (**f**), and CDRL3 (**g**) of 10LH. The CDRL loops are shown in green sticks. The map is contoured at 0.85σ for CDRL1 and 1.0σ for CDRL2 and CDRL3.

**Extended Data Figure 2.**
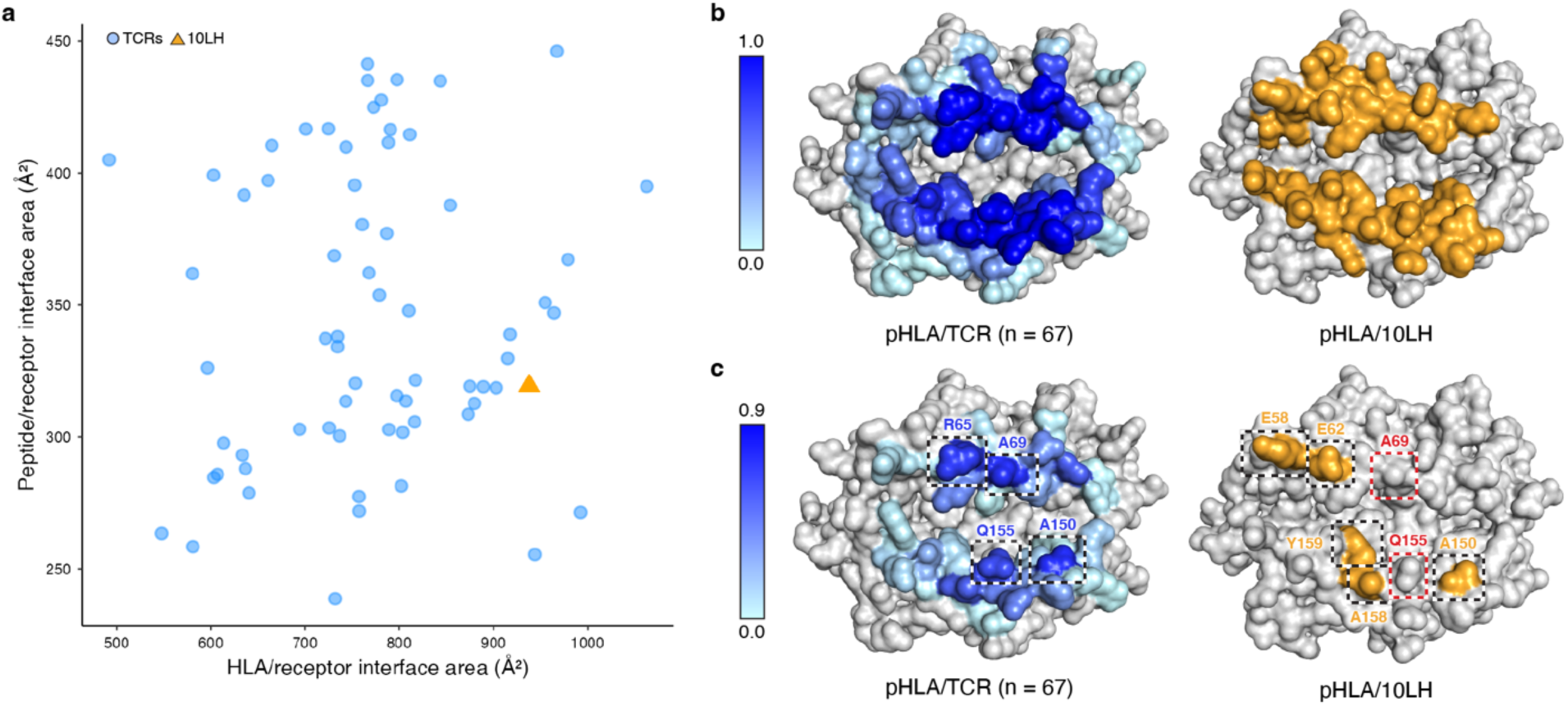
Comparison of 10LH to TCR:pHLA-I structures in the Protein Data Bank. **a**, Scatter plot depicting the interface area between the immune receptor and the HLA (x-axis) or the peptide (y-axis). The corresponding interface areas of 10LH with HLA-A*24:02 and PHOX2B are shown as an orange triangle. **b,** The pHLA interface with TCRs (blue) or 10LH (orange) mapped onto the surface of a representative HLA (TCR, PDB ID: 1AO7; 10LH, PDB ID: 8EK5). **c,** Residues with the most stabilizing effect determined by PDBePISA^23^ for TCRs (blue) or 10LH (orange) mapped onto the surface of a representative HLA.

**Extended Data Figure 3.**
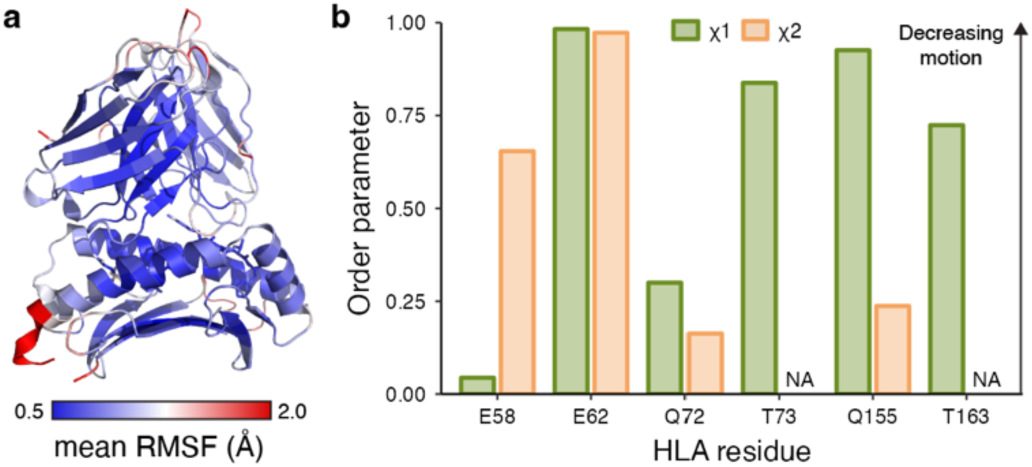
MD simulations of the ternary complex. **a**, MD simulations of 10LH bound to PHOX2B/HLA-A*24:02. HLA-A*24:02 and 10LH are shown in cartoon representation, the PHOX2B peptide is shown in stick representation. The structure is colored based on the mean RMSF value. Data are mean across n = 3 independent 1 μs runs. **b,** Order parameters of the χ_1_ and χ_2_ side chain torsion angles from the MD simulations of the 10LH:PHOX2B/HLA-A*24:02 complex. Data are shown as a sum of n = 3 independent 1 µs MD runs. NA: χ_2_ torsion angle not applicable.

**Extended Data Figure 4.**
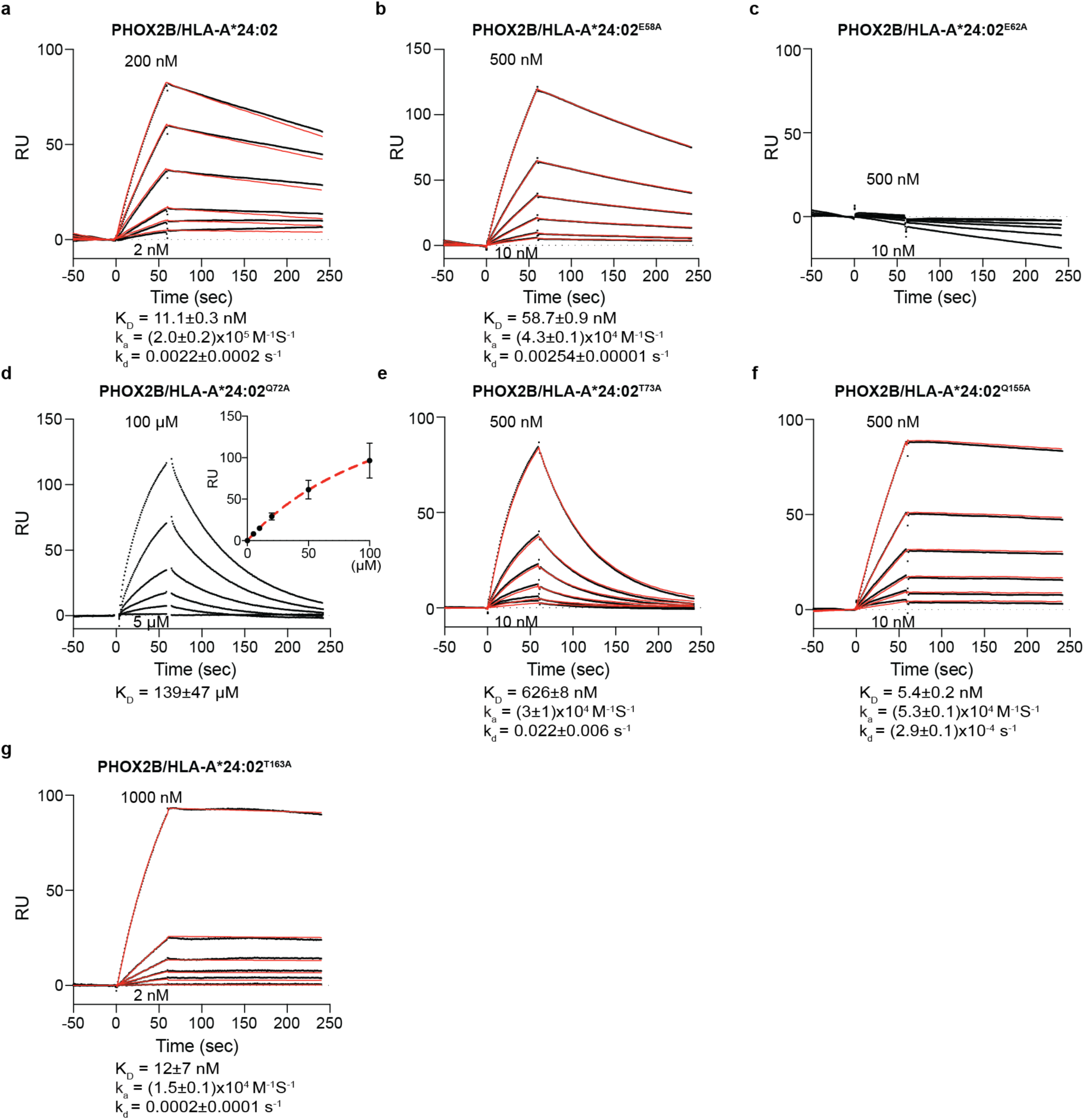
Direct interaction between 10LH and alanine-scanned HLA-A*24:02 mutants. **a-g**, SPR sensorgrams of various concentrations of the WT (**a**), E58A (**b**), E62A (**c**), Q72A (**d**), T73A (**e**), Q155A (**f**), and T163A (**g**) HLA-A*24:02 mutants flowed over a streptavidin chip coupled with 10LH-biotin. The concentrations of analyte for the top and the bottom sensorgrams are noted. Data are mean ± SD, where n = 3 technical replicates. Fits from the kinetic analysis are shown with red lines. K_D_, equilibrium constant; k_a_, association rate constant; k_d_, dissociation rate constant; RU, resonance units.

**Extended Data Figure 5.**
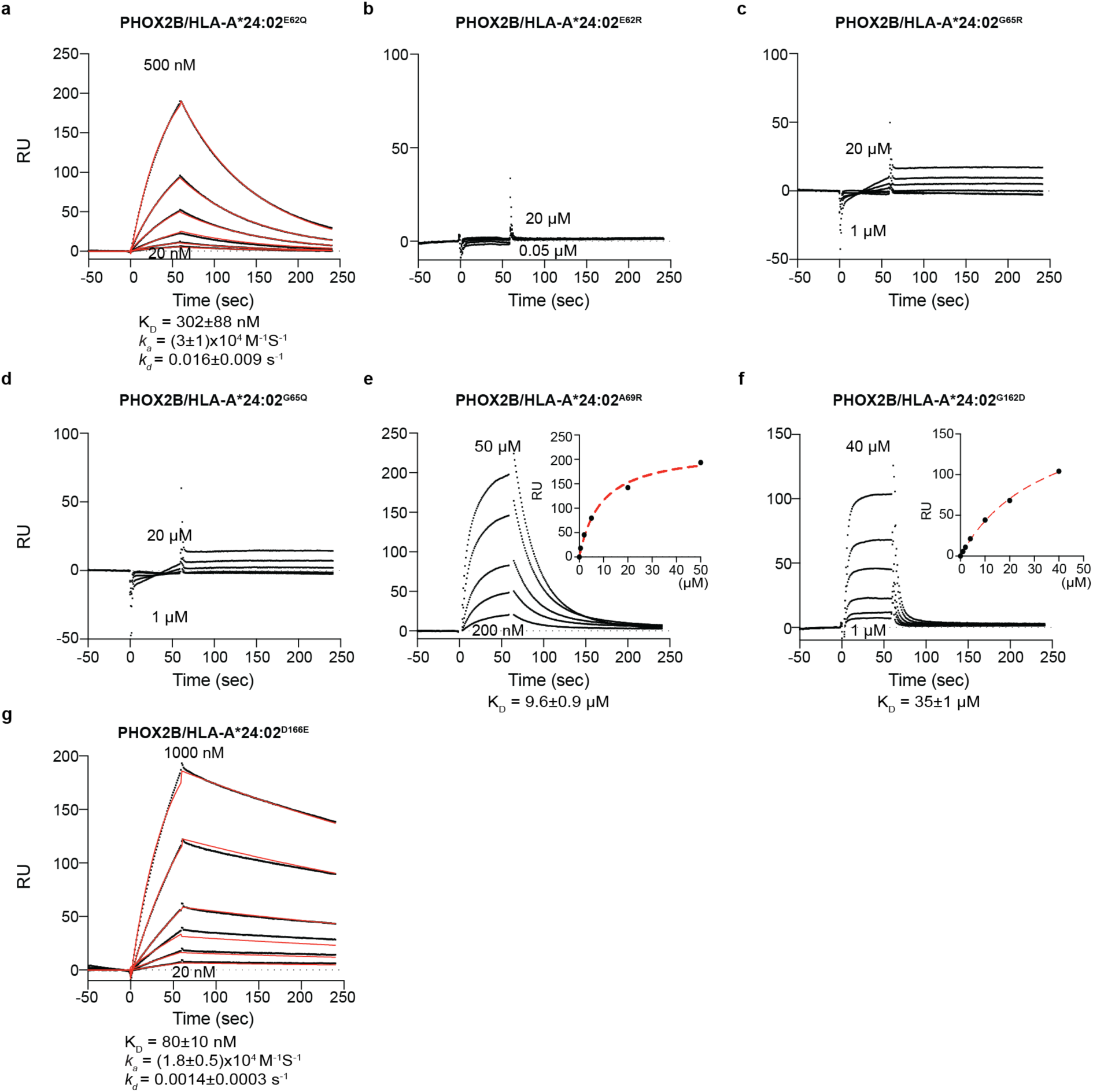
Direct interaction between 10LH and polymorphism-scanned HLA-A*24:02 variants. **a-g**, SPR sensorgrams of various concentrations of the E62Q (**a**), E62R (**b**), G65R (**c**), G65Q (**d**), A69R (**e**), G162D (**f**), and D166E (**g**) HLA-A*24:02 mutants flowed over a streptavidin chip coupled with 10LH-biotin. The concentrations of analyte for the top and the bottom sensorgrams are noted. Data are mean ± SD, where n = 3 technical replicates. Fits from the kinetic analysis are shown with red lines. K_D_, equilibrium constant; k_a_, association rate constant; k_d_, dissociation rate constant; RU, resonance units.

**Extended Data Figure 6.**
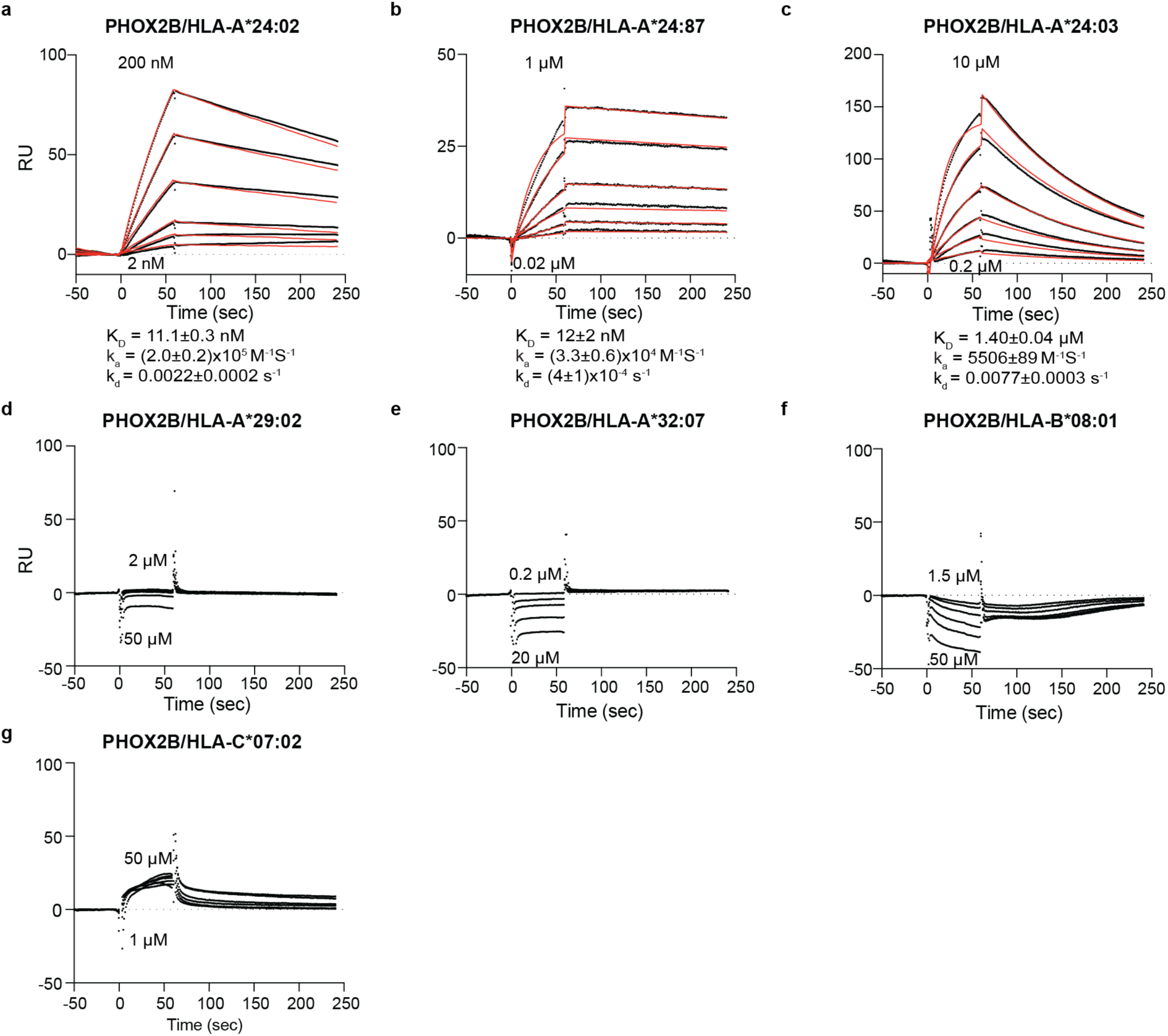
Direct interaction between 10LH and predicted HLA allotypes. **a-g**, SPR sensorgrams of various concentrations of HLA-A*24:02 (**a**), HLA-A*24:87 (**b**), HLA-A*24:03 (**c**), HLA-A*29:02 (**d**), HLA-A*32:07 (**e**), HLA-B*08:01 (**f**), and HLA-C*07:02 (**g**) flowed over a streptavidin chip coupled with 10LH-biotin. The concentrations of analyte for the top and the bottom sensorgrams are noted. Data are mean ± SD, where n = 3 technical replicates. Fits from the kinetic analysis are shown with red lines. K_D_, equilibrium constant; k_a_, association rate constant; k_d_, dissociation rate constant; RU, resonance units.

**Extended Data Figure 7.**
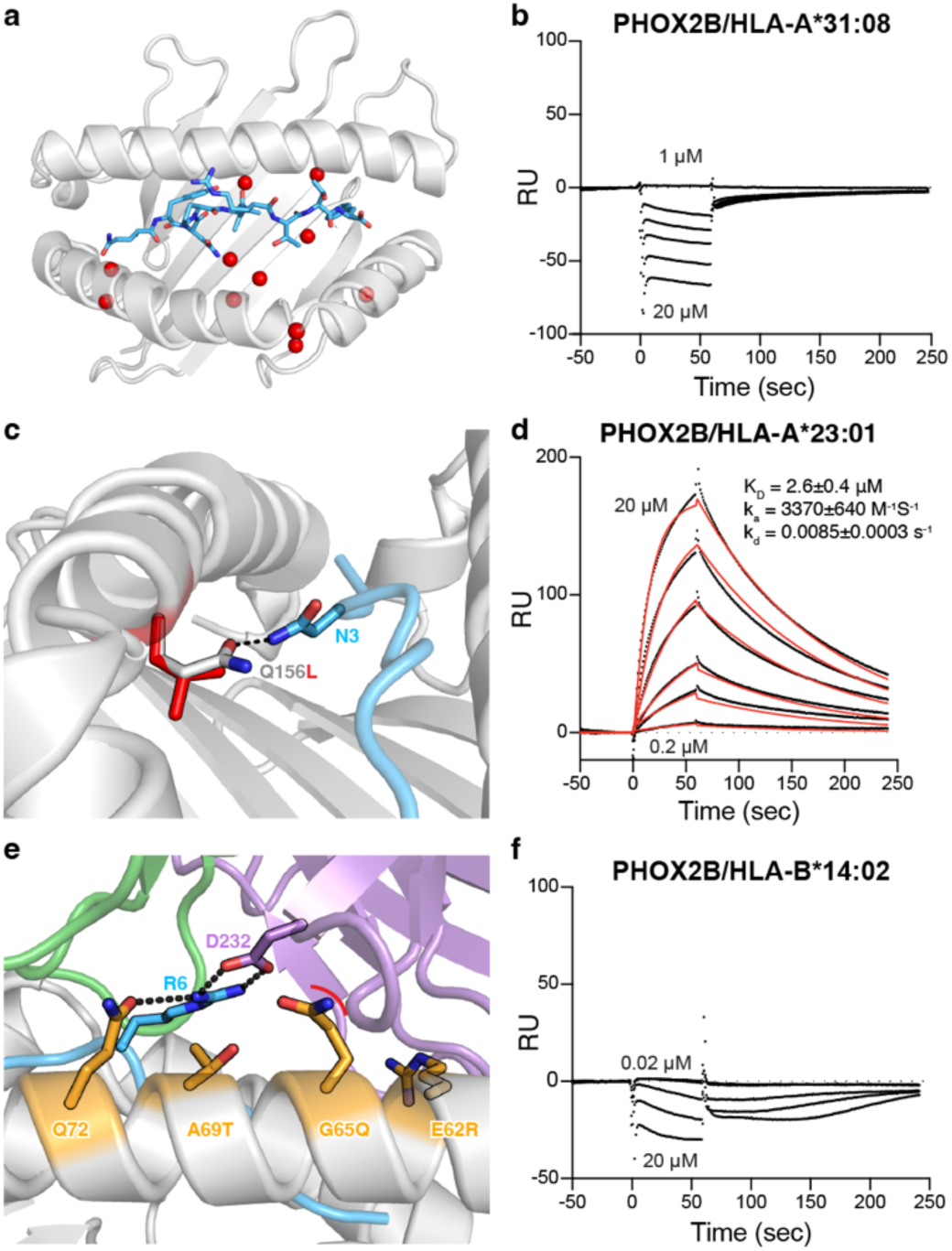
Structural basis of 10LH-specific HLA allelic restriction. **a**, Polymorphisms (red spheres) present in HLA-A*31:08 relative to HLA-A*24:02 mapped onto the 10LH:PHOX2B/HLA-A*24:02/β_2_m crystal structure (PDB ID 8EK5). **b,** Representative SPR sensorgram of HLA-A*31:08. **c,** Rosetta model of PHOX2B/HLA-A*23:01. HLA is shown in grey with the groove polymorphism (Q156L) in HLA-A*23:01 shown in red. The interacting side chain of the PHOX2B peptide is shown in blue. The hydrogen bond is shown as a black dashed line. **d,** Representative SPR sensorgram of HLA-A*23:01. **e,** Rosetta model of 10LH:PHOX2B/HLA-B*14:02/β_2_m. HLA framework residues are shown in orange and the heavy and light chains of 10LH are shown in purple and green, respectively. Steric clash is shown as a red curve. Hydrogen bonds are shown as black dashed lines. **f,** Representative SPR sensorgram of HLA-B*14:02. The concentrations of analyte for the top and the bottom sensorgrams are noted. Data are mean ± SD, where n = 3 technical replicates. Fits from the kinetic analysis are shown with red lines. K_D_, equilibrium constant; k_a_, association rate constant; k_d_, dissociation rate constant; RU, resonance units.

**Extended Data Figure 8.**
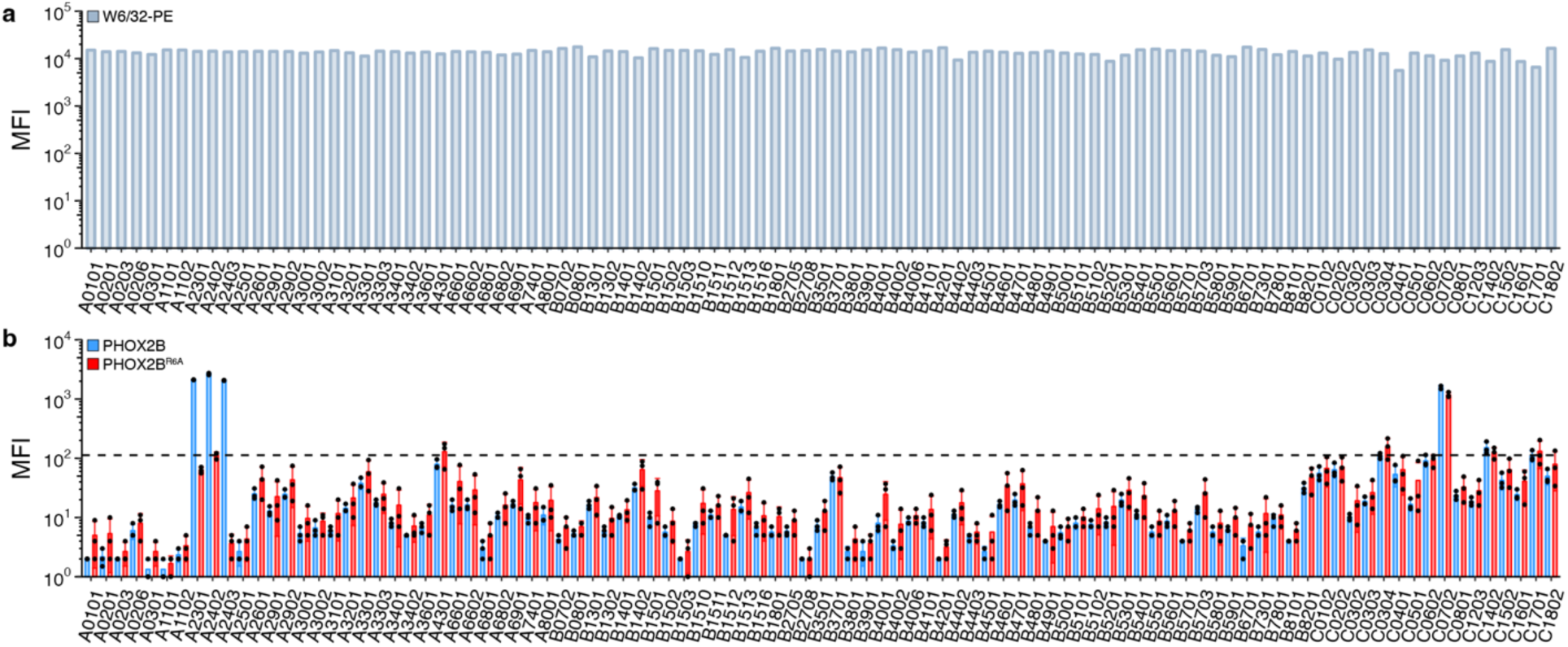
Binding levels of 10LH on HLA single antigen beads. **a**, Levels of folded HLA-I molecules captured on the SABs using the PE-conjugated W6/32 antibody (Biolegend, 311406) upon over weekend (o/w) incubation with PBS buffer, pH 7.2. Similar levels of peptide-loaded HLA-I molecules were observed across different HLA allotypes. **b,** Binding levels of 10LH tetramers at 80 nM to the beads, upon o/w incubation with excess of PHOX2B wild-type or R6A mutant (PHOX2B^R6A^) peptides. The dashed line at an MFI of 112.67 represents the threshold for non-specific binding based on the mean MFI of pre-incubation with the negative peptide (PHOX2B^R6A^) for HLA-A*24:02. Plotted data are mean ± SD from n = 3 experiments.

**Extended Data Figure 9.**
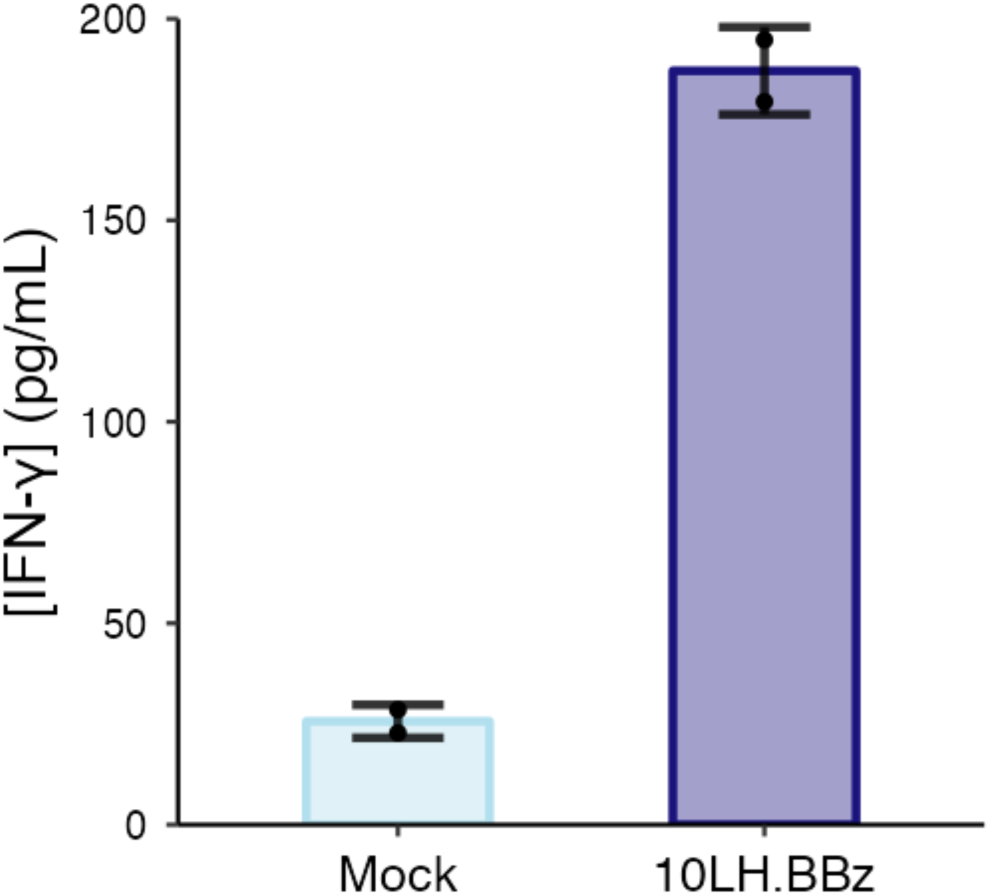
Cytokine levels of 10LH CAR T cells co-cultured with an HLA-A*24:03 neuroblastoma cell line pulsed with PHOX2B peptide. IFN-γ levels in supernatant of mock T cells (non-transduced; light blue) or 10LH.BBz CAR T cells (purple) co-cultured with HLA-A*24:03 neuroblastoma cell line SK-N-DZ pulsed with PHOX2B peptide. Exogenous peptide was added to SK-N-DZ target cells (15 µM final concentration). After four hours of incubation, 10LH.BBz CAR T cells were added to target cells at a 3:1 ET ratio. Supernatant was collected 24 hours post effector cell addition and IFN-γ levels were measured by ELISA. Values represent mean ± SD using effector cells from n = 2 biological donors, in triplicates.

**Extended Data Figure 10.**
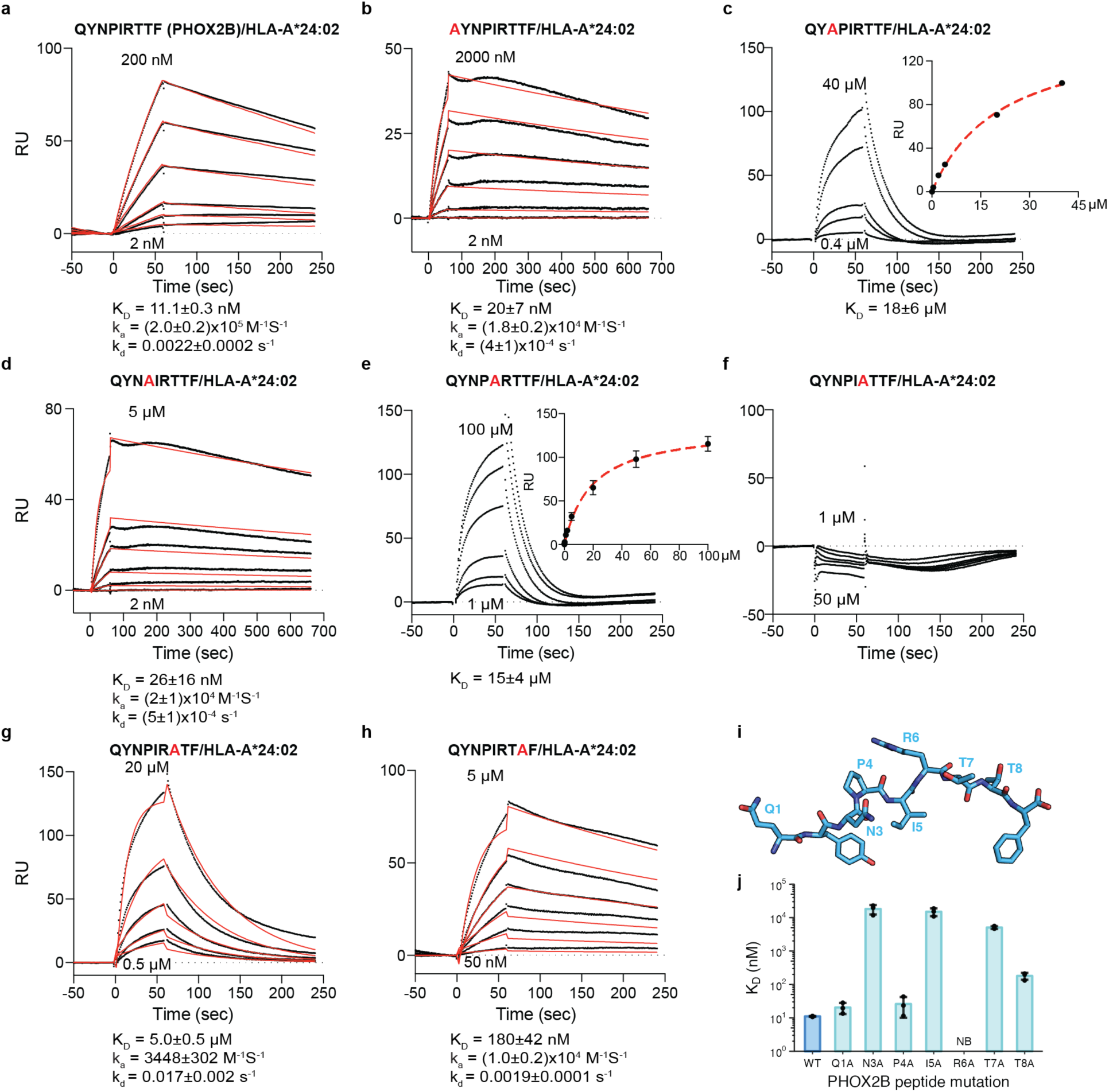
Direct interaction between 10LH and HLA-A*24:02 loaded with alanine scanned PHOX2B peptides. **a-h**, SPR sensorgrams of various concentrations of HLA-A*24:02 loaded with PHOX2B (**a**), PHOX2B^Q1A^ (**b**), PHOX2B^N3A^ (**c**), PHOX2B^P4A^ (**d**), PHOX2B^I5A^ (**e**), PHOX2B^R6A^ (**f**), PHOX2B^T7A^ (**g**), and PHOX2B^T8A^ (**h**) flowed over a streptavidin chip coupled with 10LH-biotin. The concentrations of analyte for the top and the bottom sensorgrams are noted. Data are mean ± SD, where n = 3 technical replicates. Fits from the kinetic analysis are shown with red lines. K_D_, equilibrium constant; k_a_, association rate constant; k_d_, dissociation rate constant; RU, resonance units. **i,** Stick representation of the PHOX2B peptide (PDB ID 8EK5) with non-anchor residues labeled. **j,** SPR determined K_D_ values for an alanine scan of the PHOX2B peptide. Data are mean ± SD for n = 3 technical replicates.

**Extended Data Figure 11.**
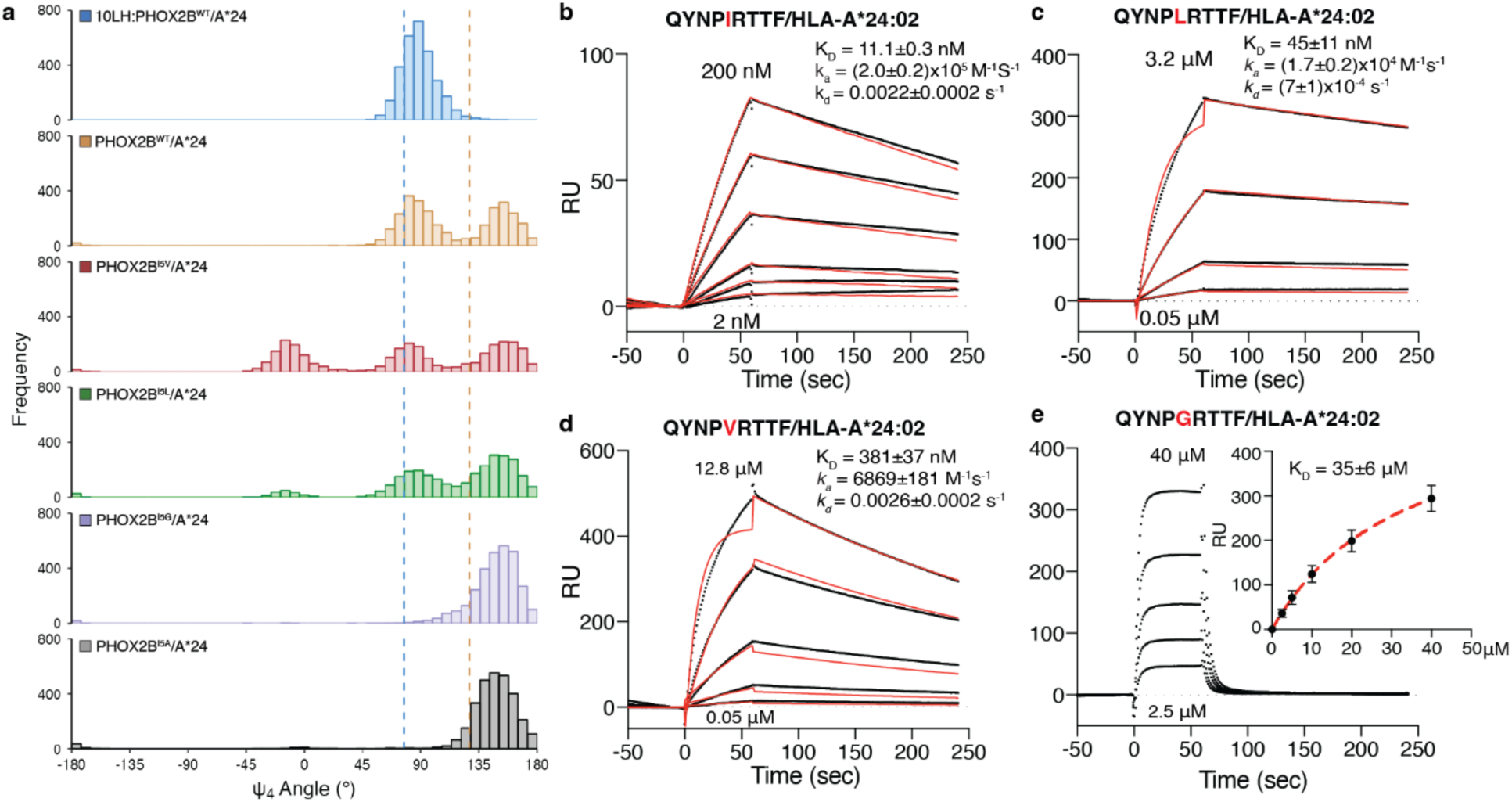
Direct interaction between 10LH and HLA-A*24:02 loaded with I5 scanned PHOX2B peptides. **a**, Histogram depicting the frequency of Ψ_4_ backbone dihedral angle values for MD simulations of the bound (blue) and unbound (brown) PHOX2B/HLA-A*24:02 structures as well as the PHOX2B^I5V^/HLA-A*24:02 (red), PHOX2B^I5L^/HLA-A*24:02 (green), PHOX2B^I5G^/HLA-A*24:02 (purple), and PHOX2B^I5A^/HLA-A*24:02 (black) variants. The dashed lines correspond to the values in the crystal structures of 10LH:PHOX2B/HLA-A*24:02/β_2_m (blue) and PHOX2B/HLA-A*24:02 (brown). Data represent 3000 equally spaced frames from a sum of n = 3 independent 1 μs runs. **b-e,** SPR sensorgrams of various concentrations of HLA-A*24:02 loaded with PHOX2B I5 (**b**), I5L (**c**), I5V (**d**), and I5G (**e**) scanning peptides flowed over a streptavidin chip coupled with 10LH-biotin. The concentrations of analyte for the top and the bottom sensorgrams are noted. Data are mean ± SD, where n = 3 technical replicates. Fits from the kinetic analysis are shown with red lines. K_D_, equilibrium constant; k_a_, association rate constant; k_d_, dissociation rate constant; RU, resonance units.

**Extended Data Figure 12.**
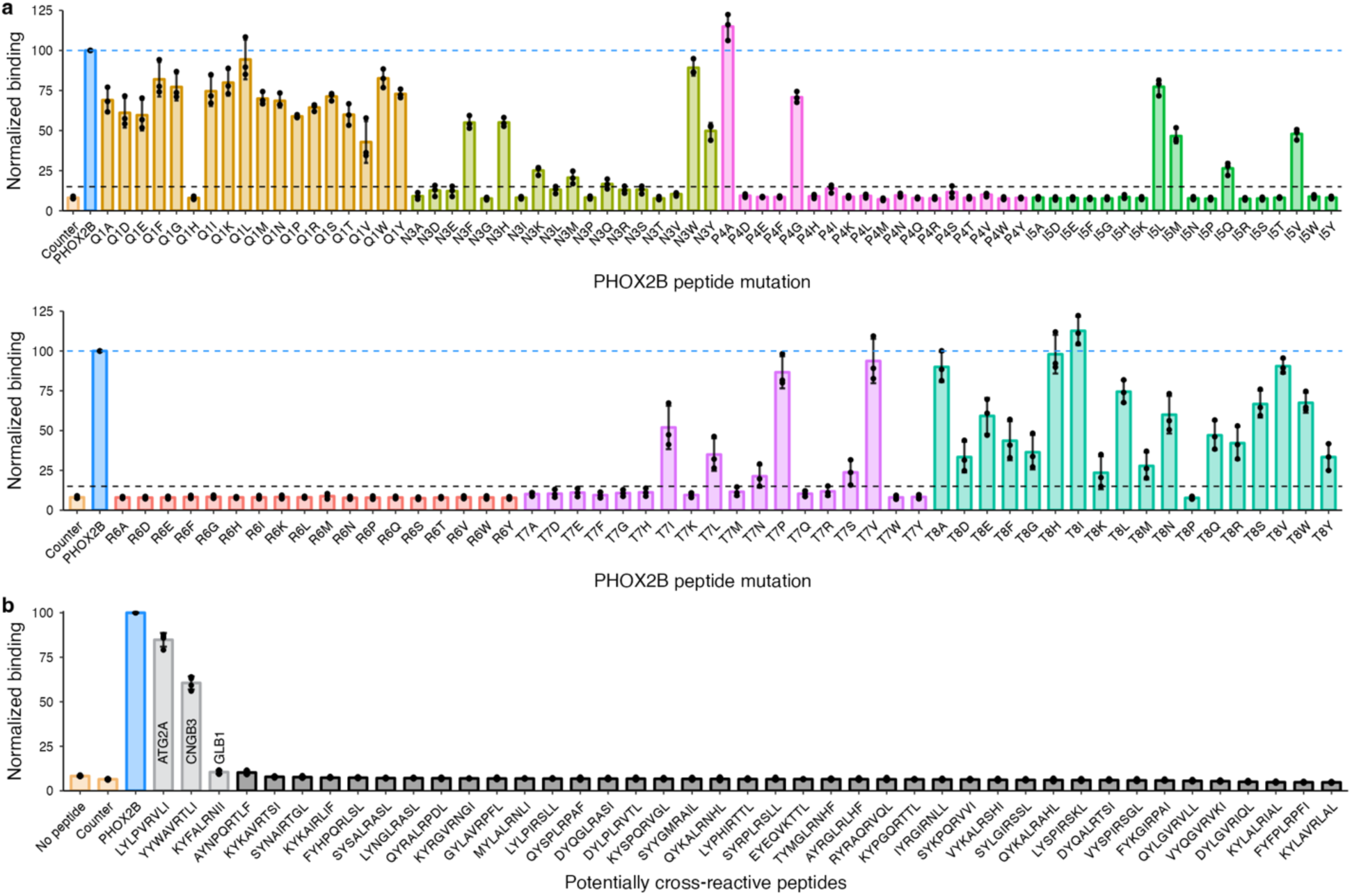
Peptide X-scan/ScanProsite analysis. **a**, Bead binding assay with 10LH scFv and PHOX2B X-scan peptides. Each non-anchor position of the PHOX2B peptide was substituted with every other amino acid except cysteine. Normalized binding expressed as mean fluorescence (488 nm) of the percent of PHOX2B peptide. The mean fluorescence for the PHOX2B peptide is shown as a blue dashed line and a black dashed line is shown at a normalized binding value of 15%, denoting the threshold for deeming an amino acid as tolerated. Values represent mean ± SD for n = 3. **b**, Bead binding assay with the 10LH scFv and potential cross-reactive peptides presented on HLA-A*24:02. Peptide sequences were generated from the X-scan/ScanProsite analysis that match sequences found in the normal human proteome. Normalized binding expressed as fluorescence (488 nm) of the percent of PHOX2B peptide.

**Extended Data Figure 13.**
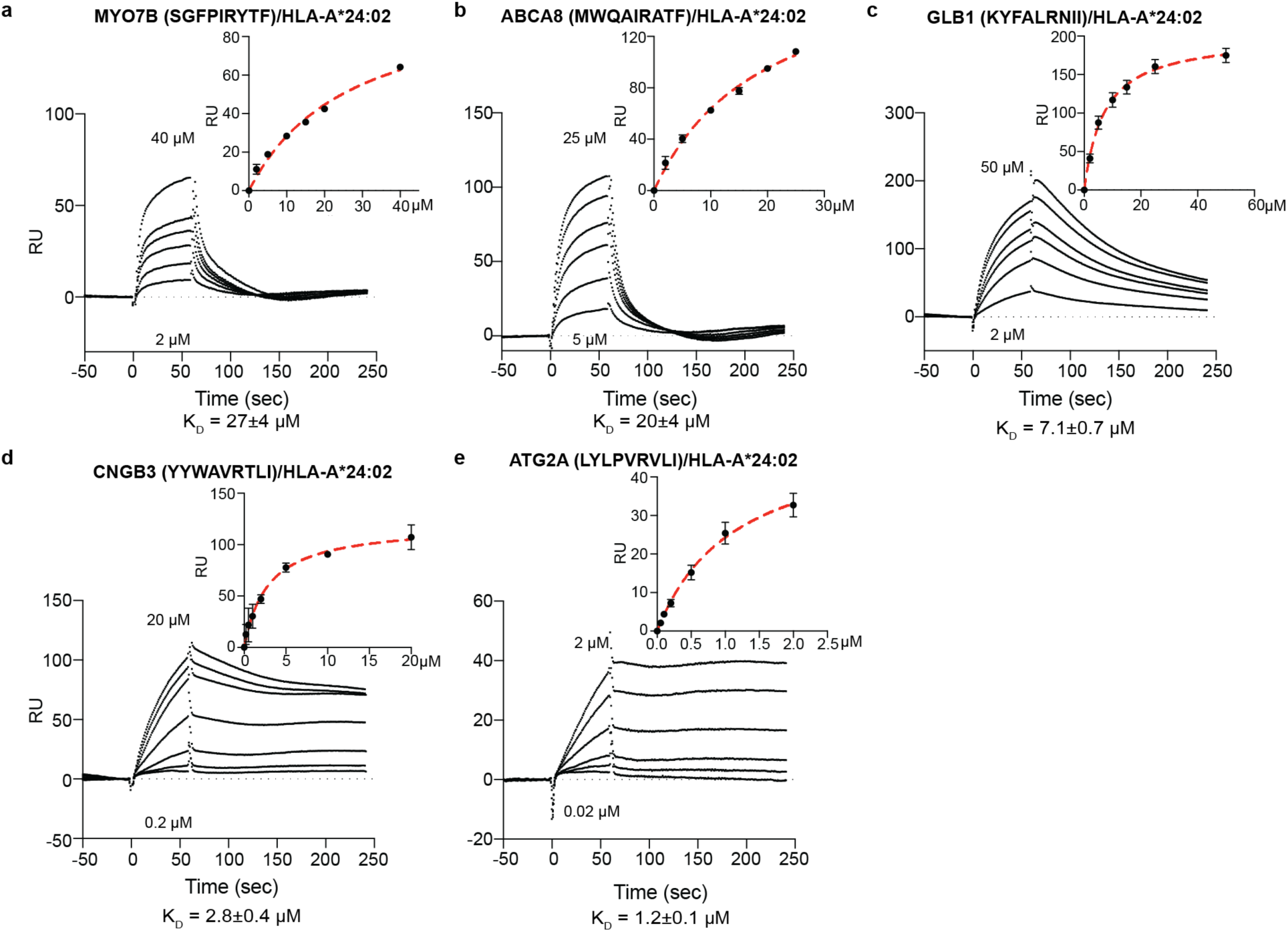
Direct interaction between 10LH and cross-reactive peptide-loaded HLA-A*24:02. **a-e**, SPR sensorgrams of various concentrations of HLA-A*24:02 loaded with MYO7B SGFPIRYTF (**a**) ABCA8 MWQAIRATF (**b**), GLB1 KYFALRNII (**c**), CNGB3 YYWAVRTLI (**d**), and ATG2A LYLPVRVLI (**e**) flowed over a streptavidin chip coupled with 10LH-biotin. The concentrations of analyte for the top and the bottom sensorgrams are noted. Data are mean ± SD, where n = 2 technical replicates. Fits from the kinetic analysis are shown with red lines. K_D_, equilibrium constant; k_a_, association rate constant; k_d_, dissociation rate constant; RU, resonance units.

**Extended Data Figure 14.**
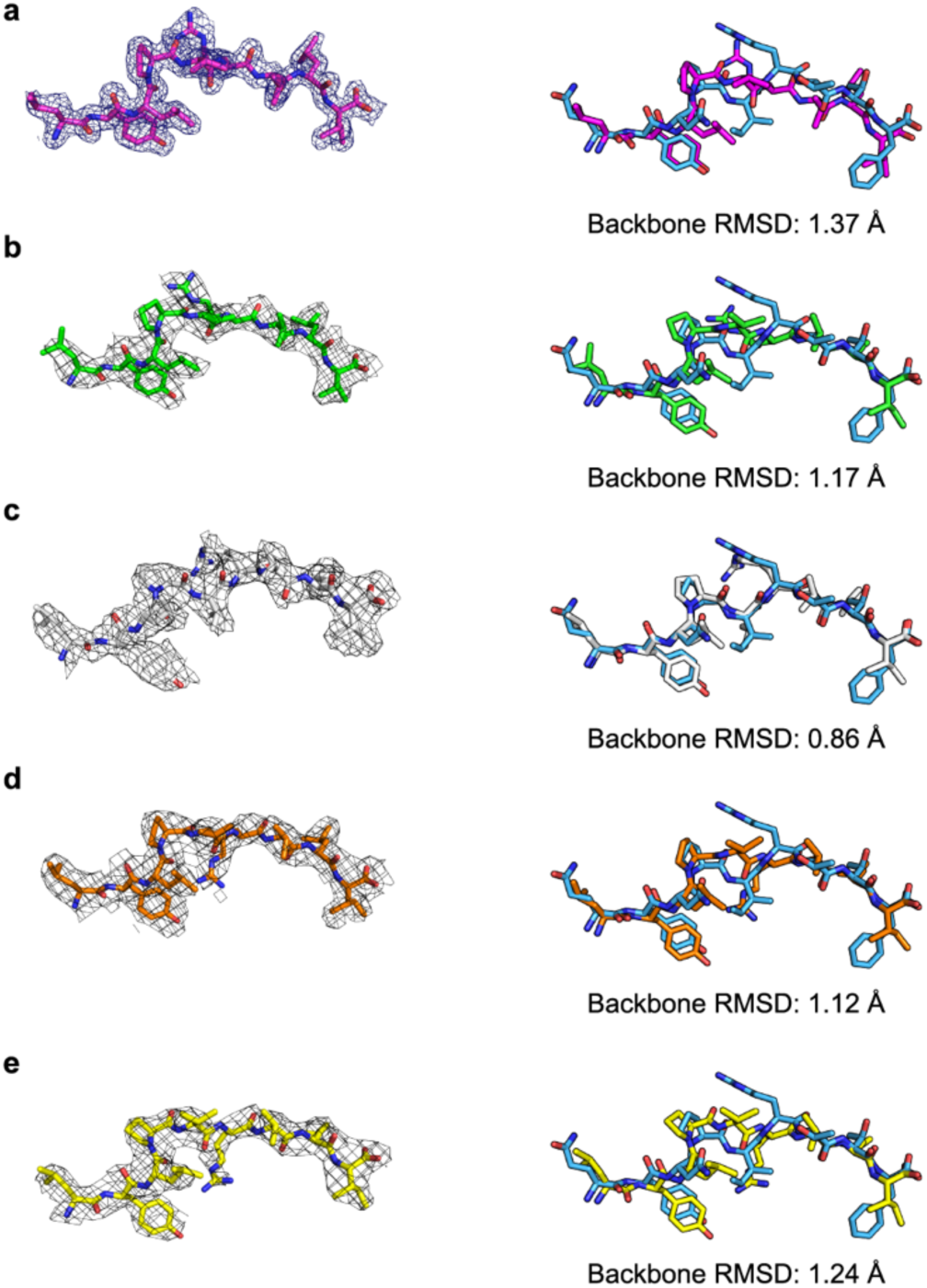
Structure determination of the ATG2A/HLA-A*24:02 complex. **a**, 2Fo-Fc omit map (black mesh) around the ATG2A peptide bound to HLA-A*24:02 contoured at 1.1σ; structure solved at 1.8 Å. To the right, a superimposition of this peptide backbone (magenta) with the peptide backbone from the ternary complex (blue) via the peptide backbone heavy atoms; RMSD is depicted below the structures. **b-e,** 2Fo-Fc omit map (black mesh) around the ATG2A peptide bound to HLA-A*24:02 contoured at 0.75σ (**b**), 0.60σ (**c**), 0.60σ (**d**), and 0.85σ (**e**); structure solved at 3.0 Å. To the right is a superimposition of the peptide backbone with the peptide backbone from the ternary complex (blue) via the peptide backbone heavy atoms; RMSD is depicted below the structures.

**Extended Data Figure 15.**
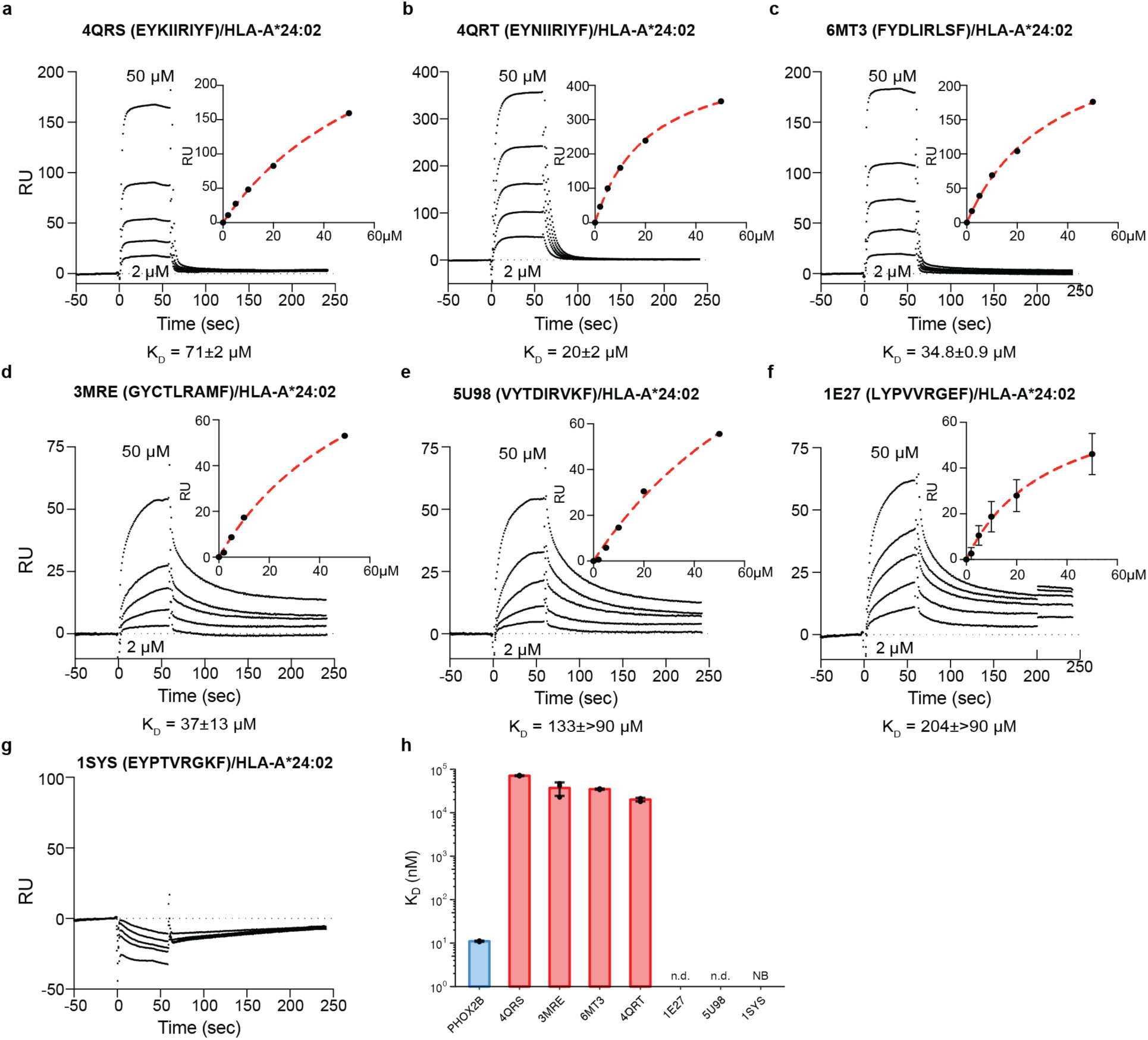
Direct interaction between 10LH and structural-guided cross-reactive peptide-loaded HLA-A*24:02. **a-g**, SPR sensorgrams of various concentrations of HLA-A*24:02 loaded with 4QRS (EYKIIRIYF) (**a**), 4QRT (EYNIIRIYF) (**b**), 6MT3 (FYDLIRLSF) (**c**), 3MRE (GYCTRLAMF) (**d**), 5U98 (VYTDIRKVF) (**e**), 1E27 (LYPVVRGEF) (**f**), and 1SYS (EYPTVRGKF) (**g**) flowed over a streptavidin chip coupled with 10LH-biotin. The concentrations of analyte for the top and the bottom sensorgrams are noted. Fits from the kinetic analysis are shown with red lines. K_D_, equilibrium constant; k_a_, association rate constant; k_d_, dissociation rate constant; RU, resonance units. **h,** SPR determined K_D_ values for structure-guided cross-reactive peptides. Data are mean ± SD for n = 3 technical replicates.

**Extended Data Table 1.** Crystallography data collection and refinement statistics.

|  | 10LH:PHOX2B/HLA-A*24:02/β<sub>2</sub>m | ATG2A/HLA-A*24:02 | ATG2A/HLA-A*24:02 |
| --- | --- | --- | --- |
| Data collection |  |  |  |
| Space group | P 1 21 1 | I 2 3 | P1 21 1 |
| Cell dimensions |  |  |  |
| a, b, c (Å) | 47.09, 174.56, 47.19 | 155.6, 155.6, 155.6 | 109.51, 76.19, 111.77 |
| α, β, γ (°) | 90.0, 118.75, 90.0 | 90.0, 90.0, 90.0 | 90.0, 109.7, 90.00 |
| Resolution (Å) * | 2.11 (2.14-2.108) | 1.8 (1.84-1.8) | 3.0 (3.21-3.05) |
| Rmerge | 0.095 (0.638) | 0.22(2.9) | 0.388 (1.20) |
| I/σ(I) | 4.5 (2.4) | 10.7 (1.2) | 2.8 (0.9) |
| Completeness (%) | 97.4 (97.1) | 100 (100) | 99.8 (100.0) |
| Redundancy | 3.8 (3.7) | 15.2 (15.7) | 3.5 (3.6) |
| Refinement |  |  |  |
| Resolution (Å) | 43.68-2.11 | 110.07-1.8 | 105.19-3.05 |
| No. reflections | 35,362 | 57,945 | 34,967 |
| Rwork / Rfree | 18.1 / 21.2 | 19.2/21.8 | 19.3/26.9 |
| No. atoms |  |  |  |
| Protein | 4,857 | 3,144 | 12,604 |
| Ligand | 20 | 24 | 0 |
| Water | 306 | 412 | 0 |
| B-factors (Å<sup>2</sup>) | 35.17 | 30.12 | 36.83 |
| R.m.s. deviations |  |  |  |
| Bond lengths (Å) | 0.008 | 0.009 | 0.009 |
| Bond angles (°) | 1.2 | 0.92 | 1.177 |

\*Values in parentheses are for highest-resolution shell.

**Extended Data Table 2.** Expanded American population coverage of 10LH. AFA: African American; API: Asian or Pacific Islander; CAU: Caucasian; HIS: Hispanic; NAM: Native American.

| Allele | Population frequency (%) |  |  |  |  |  | Polymorphisms<br>relative to HLA-<br>A*24:02 |
| --- | --- | --- | --- | --- | --- | --- | --- |
|  | AFA | API | CAU | HIS | NAM | American<br>(total) |  |
| HLA-A*24:02 | 4.90 | 31.63 | 17.03 | 23.17 | 24.20 | 17.65 | N/A |
| HLA-A*24:03 | 0.09 | 1.12 | 0.49 | 0.61 | 0.91 | 0.51 | D166E, G167W |
| HLA-A*24:05 | 0.00 | 0.00 | 0.00 | 0.04 | 0.00 | 0.01 | None |
| HLA-A*24:07 | 0.10 | 4.43 | 0.02 | 0.06 | 0.08 | 0.31 | None |
| HLA-A*24:10 | 0.00 | 0.48 | 0.00 | 0.00 | 0.00 | 0.03 | T163R, D166E,<br>G167W |
| HLA-A*24:17 | 0.00 | 0.47 | 0.00 | 0.00 | 0.01 | 0.03 | None |
| HLA-A*24:20 | 0.00 | 0.09 | 0.00 | 0.00 | 0.00 | 0.01 | None |
| HLA-A*24:22 | 0.00 | 0.00 | 0.00 | 0.08 | 0.00 | 0.02 | Q156W, A158V,<br>D166E, G167W |
| HLA-A*24:25 | 0.00 | 0.00 | 0.00 | 0.28 | 0.02 | 0.06 | None |
| HLA-A*23:01 | 20.61 | 0.73 | 4.13 | 6.73 | 3.84 | 6.52 | H151R, Q156L |
| **10 alleles** | **25.70** | **38.95** | **21.67** | **30.97** | **29.06** | **25.15** | **N/A** |

**Extended Data Table 3.**
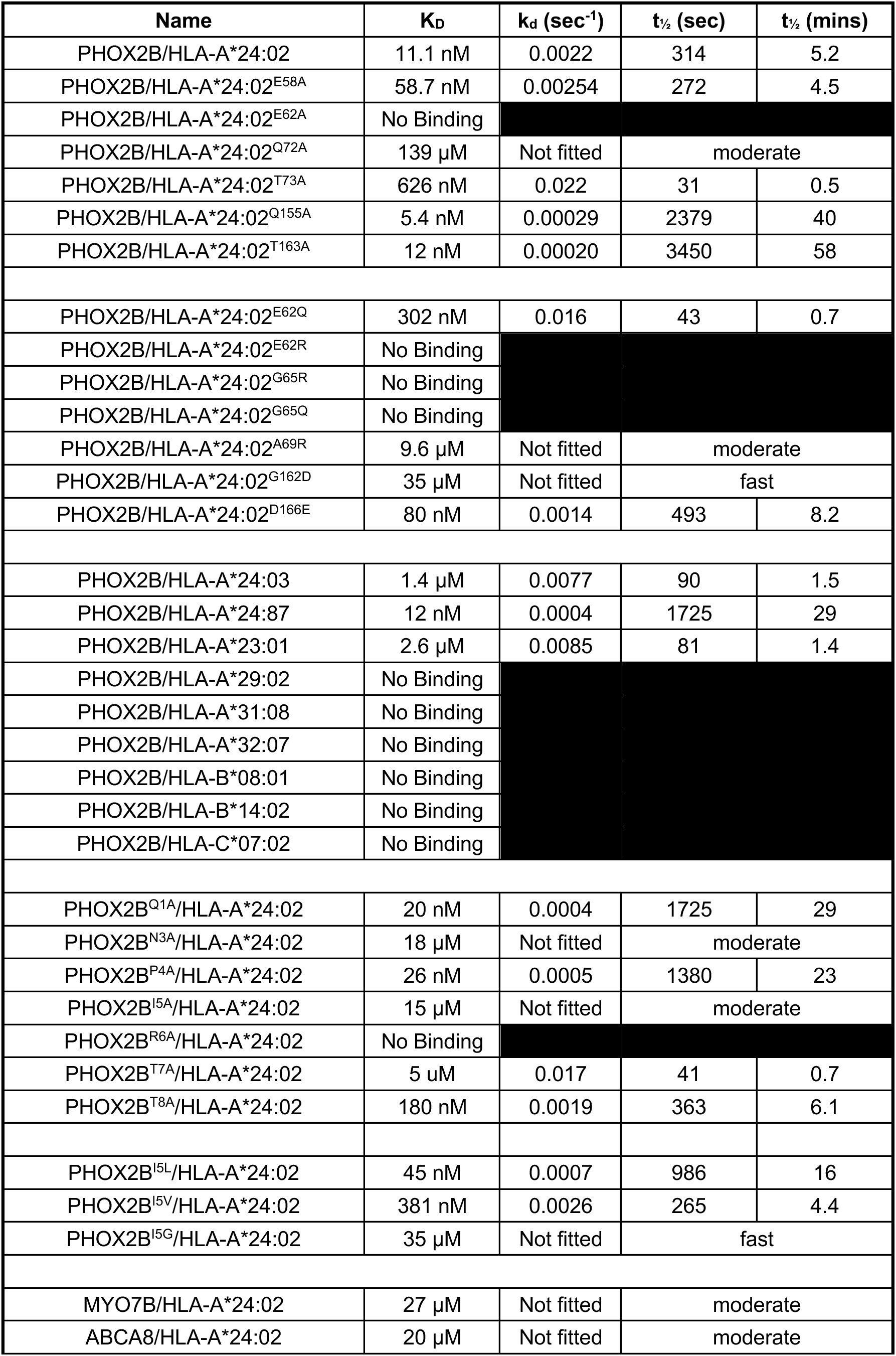
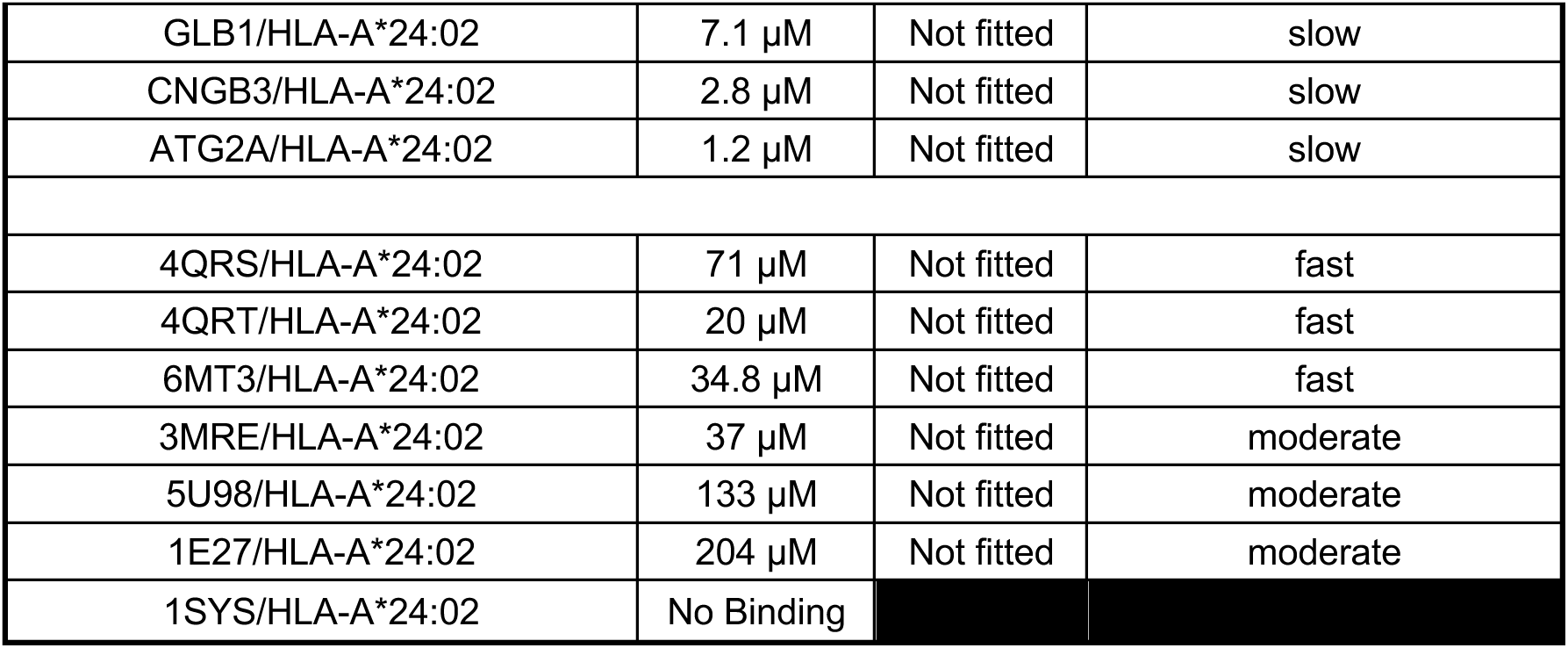
Summary of SPR parameters of K_D_, k_d_ and the 10LH complex half-life, t_1/2_. Moderate or fast t**_1/2_** indicates the k**_d_** value is too big to be fitted, beyond the instrument detection limit. Slow t**_1/2_** indicates the k**_d_** is too small to be fitted, beyond the instrument detection limit. Error (SD) can be found in extended data figures.

| Name | K<sub>D</sub> | k<sub>d</sub> (sec<sup>-1</sup>) | t<sub>½</sub> (sec) | t<sub>½</sub> (mins) |
| --- | --- | --- | --- | --- |
| PHOX2B/HLA-A*24:02 | 11.1 nM | 0.0022 | 314 | 5.2 |
| PHOX2B/HLA-A*24:02<sup>E58A</sup> | 58.7 nM | 0.00254 | 272 | 4.5 |
| PHOX2B/HLA-A*24:02<sup>E62A</sup> | No Binding |  |  |  |
| PHOX2B/HLA-A*24:02<sup>Q72A</sup> | 139 μM | Not fitted | moderate |  |
| PHOX2B/HLA-A*24:02<sup>T73A</sup> | 626 nM | 0.022 | 31 | 0.5 |
| PHOX2B/HLA-A*24:02<sup>Q155A</sup> | 5.4 nM | 0.00029 | 2379 | 40 |
| PHOX2B/HLA-A*24:02<sup>T163A</sup> | 12 nM | 0.00020 | 3450 | 58 |
| PHOX2B/HLA-A*24:02<sup>E62Q</sup> | 302 nM | 0.016 | 43 | 0.7 |
| PHOX2B/HLA-A*24:02<sup>E62R</sup> | No Binding |  |  |  |
| PHOX2B/HLA-A*24:02<sup>G65R</sup> | No Binding |  |  |  |
| PHOX2B/HLA-A*24:02<sup>G65Q</sup> | No Binding |  |  |  |
| PHOX2B/HLA-A*24:02<sup>A69R</sup> | 9.6 μM | Not fitted | moderate |  |
| PHOX2B/HLA-A*24:02<sup>G162D</sup> | 35 μM | Not fitted | fast |  |
| PHOX2B/HLA-A*24:02<sup>D166E</sup> | 80 nM | 0.0014 | 493 | 8.2 |
| PHOX2B/HLA-A*24:03 | 1.4 μM | 0.0077 | 90 | 1.5 |
| PHOX2B/HLA-A*24:87 | 12 nM | 0.0004 | 1725 | 29 |
| PHOX2B/HLA-A*23:01 | 2.6 μM | 0.0085 | 81 | 1.4 |
| PHOX2B/HLA-A*29:02 | No Binding |  |  |  |
| PHOX2B/HLA-A*31:08 | No Binding |  |  |  |
| PHOX2B/HLA-A*32:07 | No Binding |  |  |  |
| PHOX2B/HLA-B*08:01 | No Binding |  |  |  |
| PHOX2B/HLA-B*14:02 | No Binding |  |  |  |
| PHOX2B/HLA-C*07:02 | No Binding |  |  |  |
| PHOX2B<sup>Q1A</sup>/HLA-A*24:02 | 20 nM | 0.0004 | 1725 | 29 |
| PHOX2B<sup>N3A</sup>/HLA-A*24:02 | 18 μM | Not fitted | moderate |  |
| PHOX2B<sup>P4A</sup>/HLA-A*24:02 | 26 nM | 0.0005 | 1380 | 23 |
| PHOX2B<sup>I5A</sup>/HLA-A*24:02 | 15 μM | Not fitted | moderate |  |
| PHOX2B<sup>R6A</sup>/HLA-A*24:02 | No Binding |  |  |  |
| PHOX2B<sup>T7A</sup>/HLA-A*24:02 | 5 uM | 0.017 | 41 | 0.7 |
| PHOX2B<sup>T8A</sup>/HLA-A*24:02 | 180 nM | 0.0019 | 363 | 6.1 |
| PHOX2B<sup>I5L</sup>/HLA-A*24:02 | 45 nM | 0.0007 | 986 | 16 |
| PHOX2B<sup>I5V</sup>/HLA-A*24:02 | 381 nM | 0.0026 | 265 | 4.4 |
| PHOX2B<sup>I5G</sup>/HLA-A*24:02 | 35 μM | Not fitted | fast |  |
| MYO7B/HLA-A*24:02 | 27 μM | Not fitted | moderate |  |
| ABCA8/HLA-A*24:02 | 20 μM | Not fitted | moderate |  |

|  |  |  |  |
| --- | --- | --- | --- |
| GLB1/HLA-A*24:02 | 7.1 μM | Not fitted | slow |
| CNGB3/HLA-A*24:02 | 2.8 μM | Not fitted | slow |
| ATG2A/HLA-A*24:02 | 1.2 μM | Not fitted | slow |
| 4QRS/HLA-A*24:02 | 71 μM | Not fitted | fast |
| 4QRT/HLA-A*24:02 | 20 μM | Not fitted | fast |
| 6MT3/HLA-A*24:02 | 34.8 μM | Not fitted | fast |
| 3MRE/HLA-A*24:02 | 37 μM | Not fitted | moderate |
| 5U98/HLA-A*24:02 | 133 μM | Not fitted | moderate |
| 1E27/HLA-A*24:02 | 204 μM | Not fitted | moderate |
| 1SYS/HLA-A*24:02 | No Binding |  |  |

## Supplementary Information

**Supplementary Figure 1.**
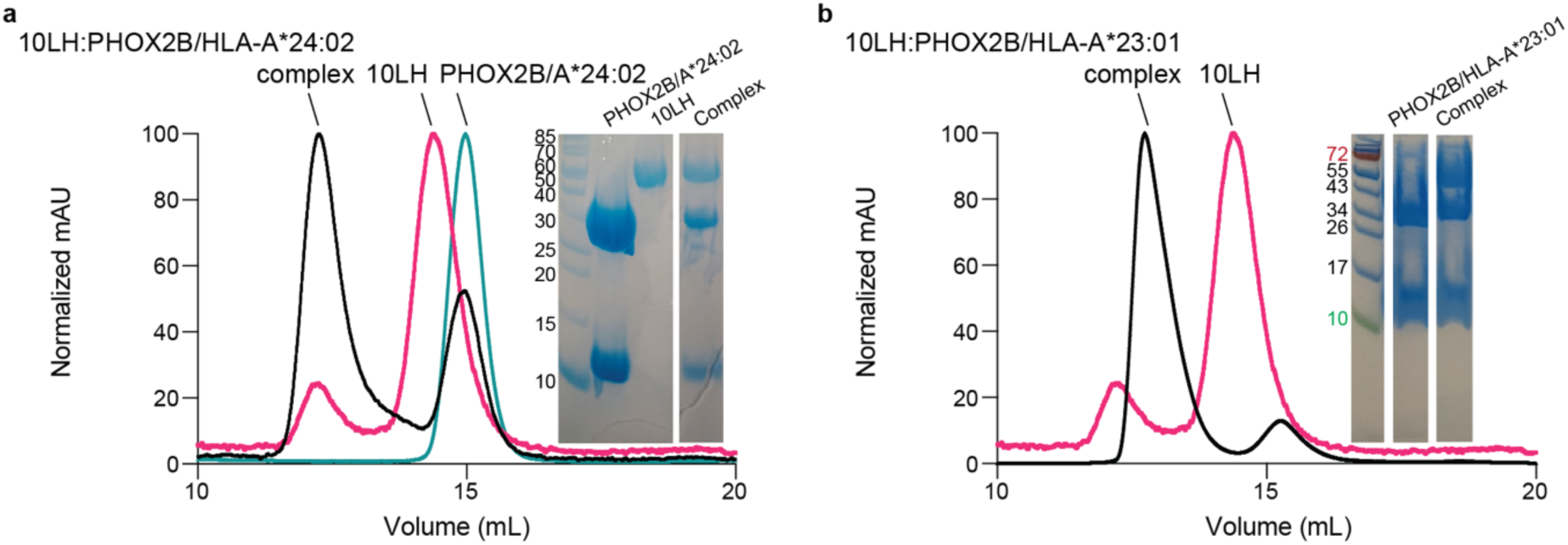
Purification of stoichiometric 10LH:PHOX2B/HLA-A*24:02/β_2_m and 10LH:PHOX2B/HLA-A*23:01/β_2_m complexes. **a-b**, Size exclusion chromatography (SEC) traces of 10LH:PHOX2B/HLA-A*24:02/β_2_m, 10LH, and PHOX2B/HLA-A*24:02/β_2_m (**a**), and 10LH:PHOX2B/HLA-A*23:01/β_2_m complex and 10LH proteins (**b**). The protein peaks were further confirmed by SDS/PAGE analysis. Gel showing bands for 10LH (∼50 kDa), HLA-A*24:02 (∼32 kDa), HLA-A*23:01 (∼32 kDa), and β_2_m (∼12 kDa).

**Supplementary Figure 2.**
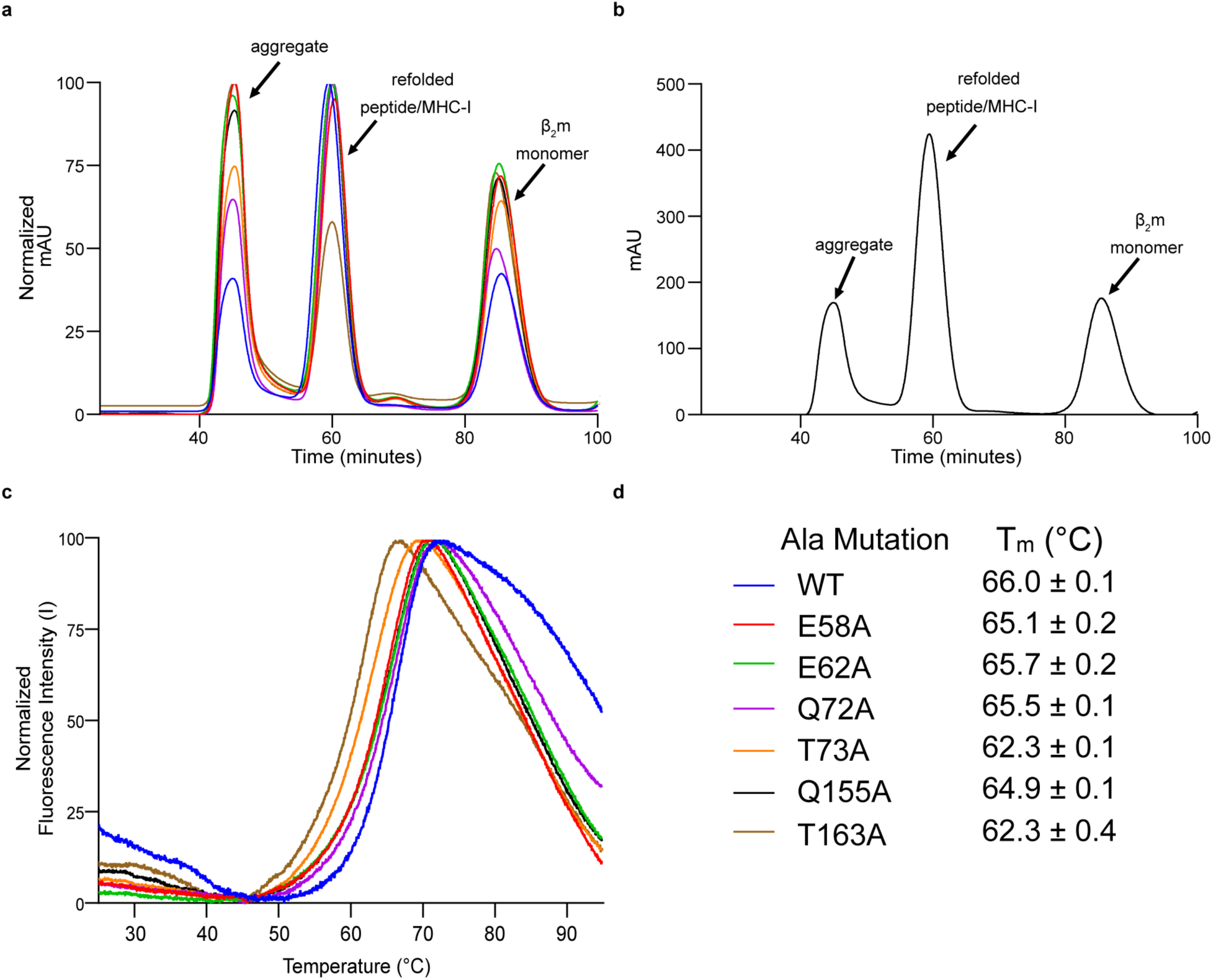
SEC and DSF data for alanine scanned HLA-A*24:02 mutants. **a**, Normalized size exclusion chromatography (SEC) traces of alanine scanned HLA-A*24:02/β_2_m refolded in the presence of PHOX2B peptide. SEC traces for alanine scanned HLA-A*24:02 molecules are color-coded, as shown in panel **d**. **b,** Representative SEC trace of PHOX2B/HLA-A*24:02/β_2_m indicating each peak identify. **c,** Normalized DSF traces of purified alanine scanned PHOX2B/HLA-A*24:02/β_2_m, color-coded as shown in panel **d**. **d,** Summary of melting temperatures (T_m_, °C) obtained from DSF experiments.

**Supplementary Figure 3.**
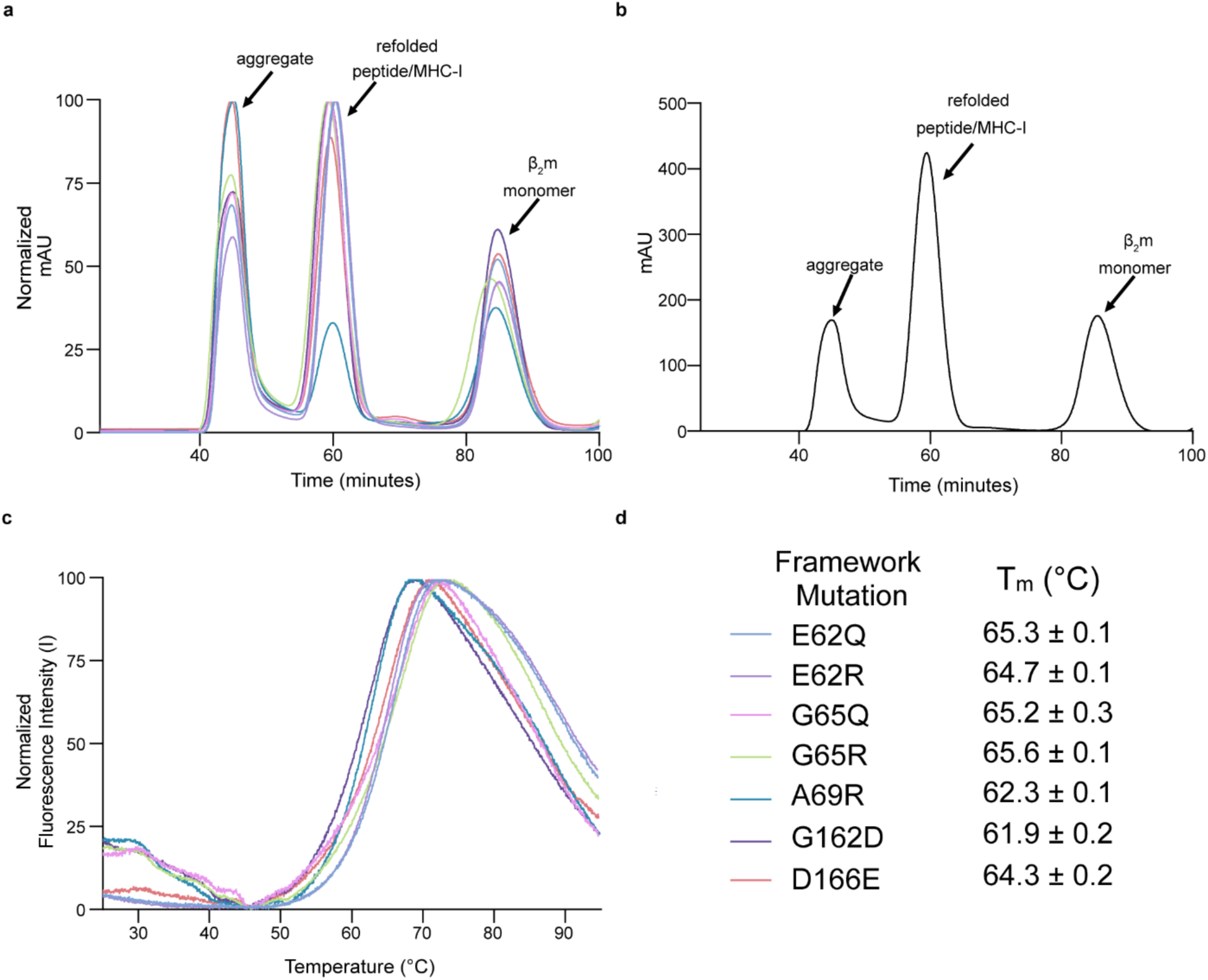
SEC and DSF data for polymorphism scanned HLA-A*24:02 variants. **a**, Normalized size exclusion chromatography (SEC) traces of polymorphism scanned HLA-A*24:02/β_2_m refolded in the presence of PHOX2B peptide. SEC traces for polymorphism scanned HLA-A*24:02 molecules are color-coded, as shown in panel **d**. **b,** Representative SEC trace of PHOX2B/HLA-A*24:02/β_2_m indicating each peak identify. **c,** Normalized DSF traces of purified polymorphism scanned PHOX2B/HLA-A*24:02/β_2_m, color-coded as shown in panel **d**. **d,** Summary of melting temperatures (T_m_, °C) obtained from DSF experiments.

**Supplementary Figure 4.**
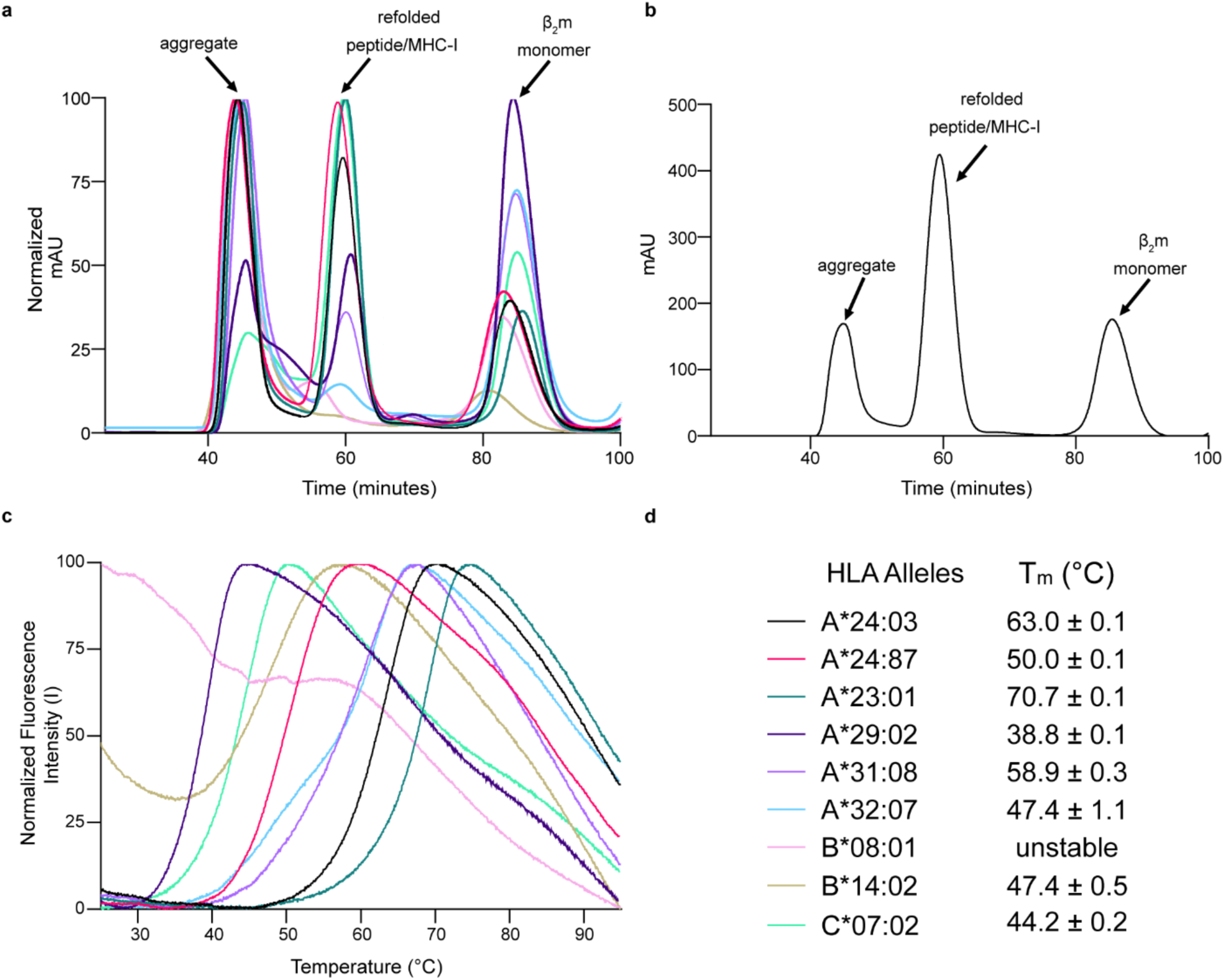
SEC and DSF data for predicted HLA allotypes. **a**, Normalized size exclusion chromatography (SEC) traces of predicted HLA allotypes refolded in the presence of PHOX2B peptide. SEC traces for predicted HLA molecules are color-coded, as shown in panel **d**. **b,** Representative SEC trace of PHOX2B/HLA-A*24:02/β_2_m indicating each peak identify. **c,** Normalized DSF traces of purified PHOX2B-loaded HLA allotypes, color-coded as shown in panel **d**. **d,** Summary of melting temperatures (T_m_, °C) obtained from DSF experiments.

**Supplementary Figure 5.**
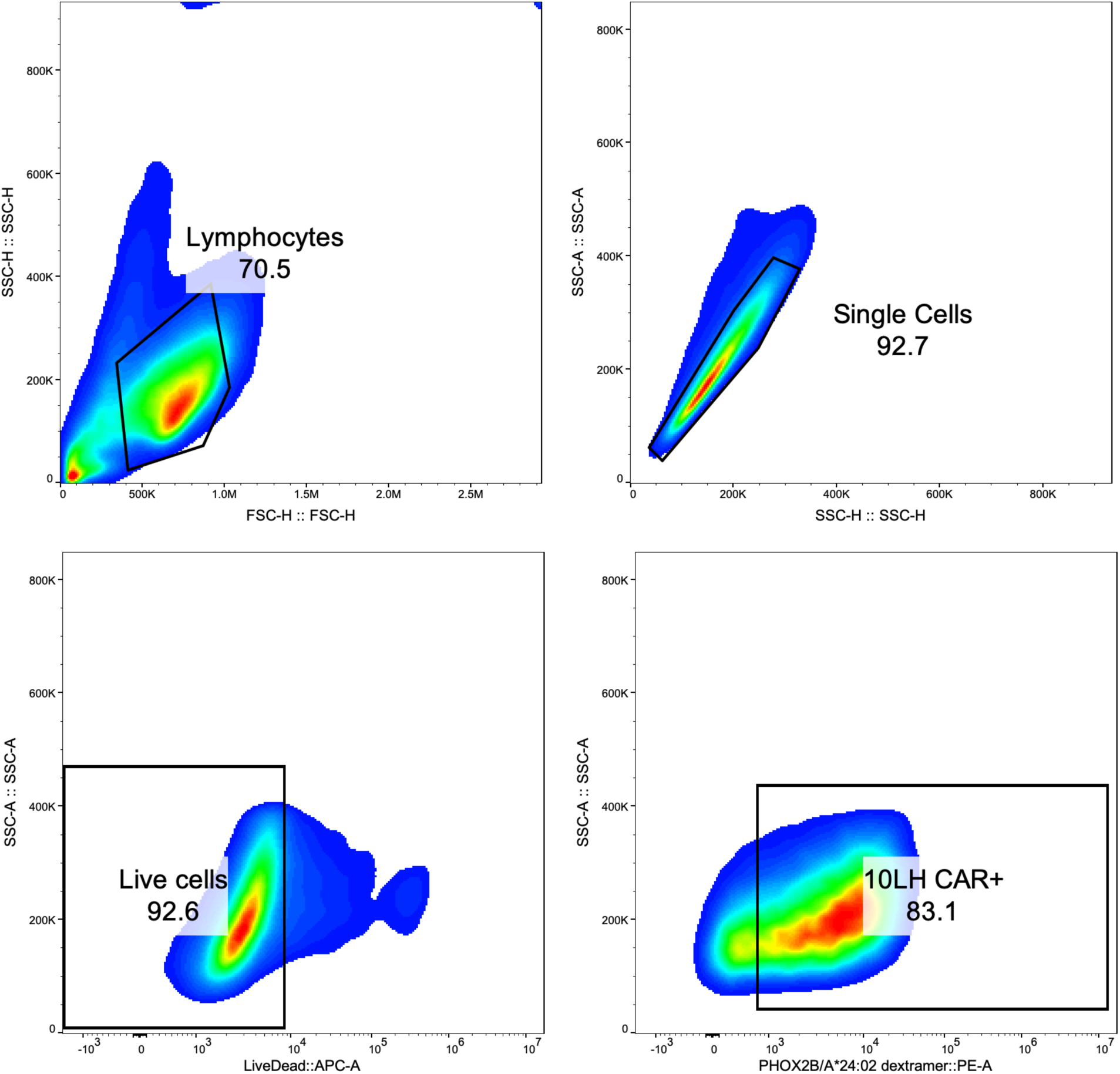
Dextramer gating strategy for verification of 10LH CAR T cell transduction efficiency. Starting event populations were gated under FSC-A/SSC-A plots for cells. T cell singlets were then identified using SSC-H/SSC-A plots. Live cells were next gated using SSC-A/APC-A plots. CAR+ cells were finally gated using SSC-A/PE-A plots.

**Supplementary Figure 6.**
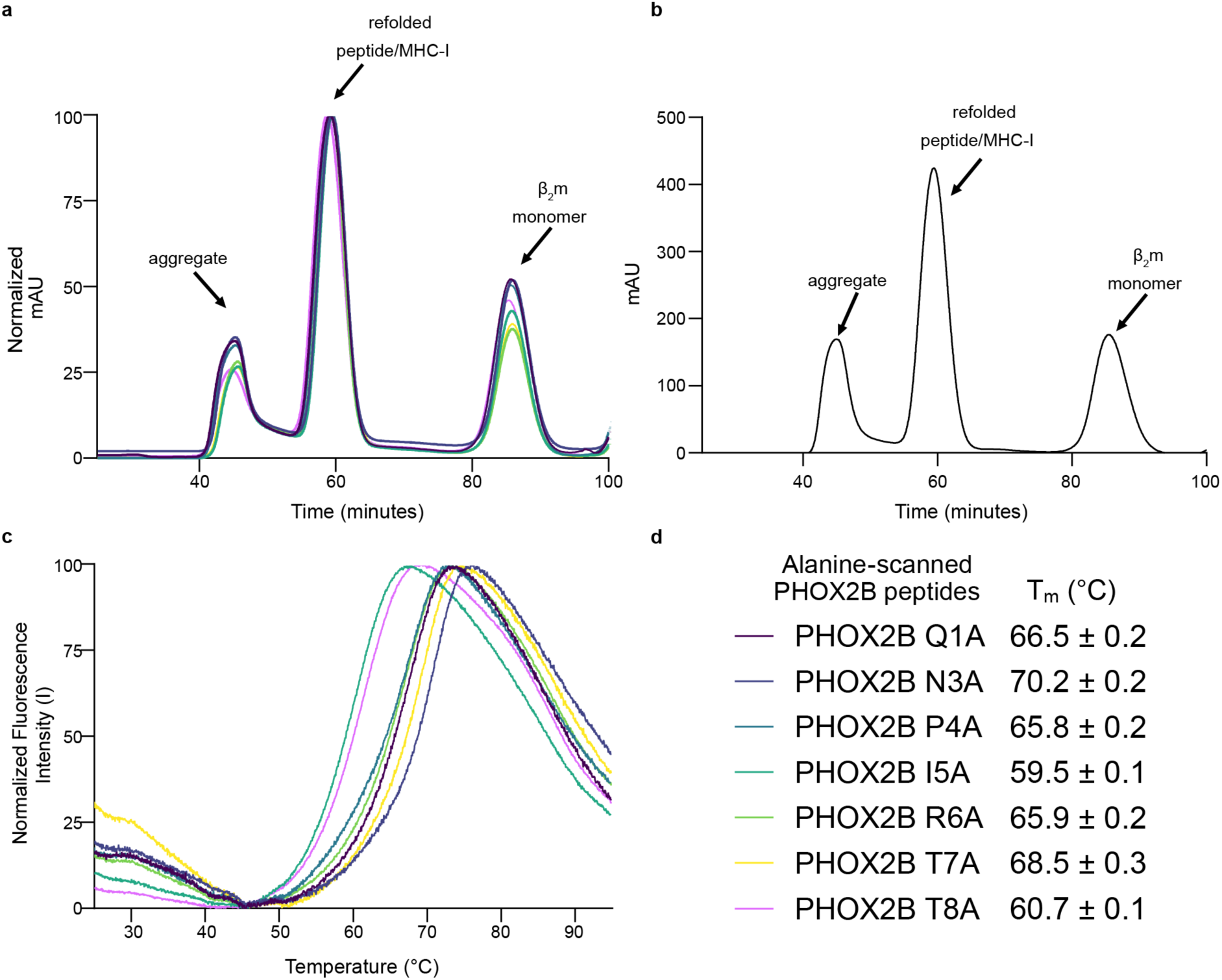
SEC and DSF data for alanine scanned PHOX2B peptides. **a**, Normalized size exclusion chromatography (SEC) traces of HLA-A*24:02/β_2_m refolded with alanine scanned PHOX2B peptides. SEC traces for predicted HLA allotypes molecules are color-coded, as shown in panel **d**. **b,** Representative SEC trace of PHOX2B/HLA-A*24:02/β_2_m indicating each peak identify. **c,** Normalized DSF traces of HLA-A*24:02/β_2_m refolded with alanine scanned PHOX2B peptides, color-coded as shown in panel **d**. **d,** Summary of melting temperatures (T_m_, °C) obtained from DSF experiments.

**Supplementary Figure 7.**
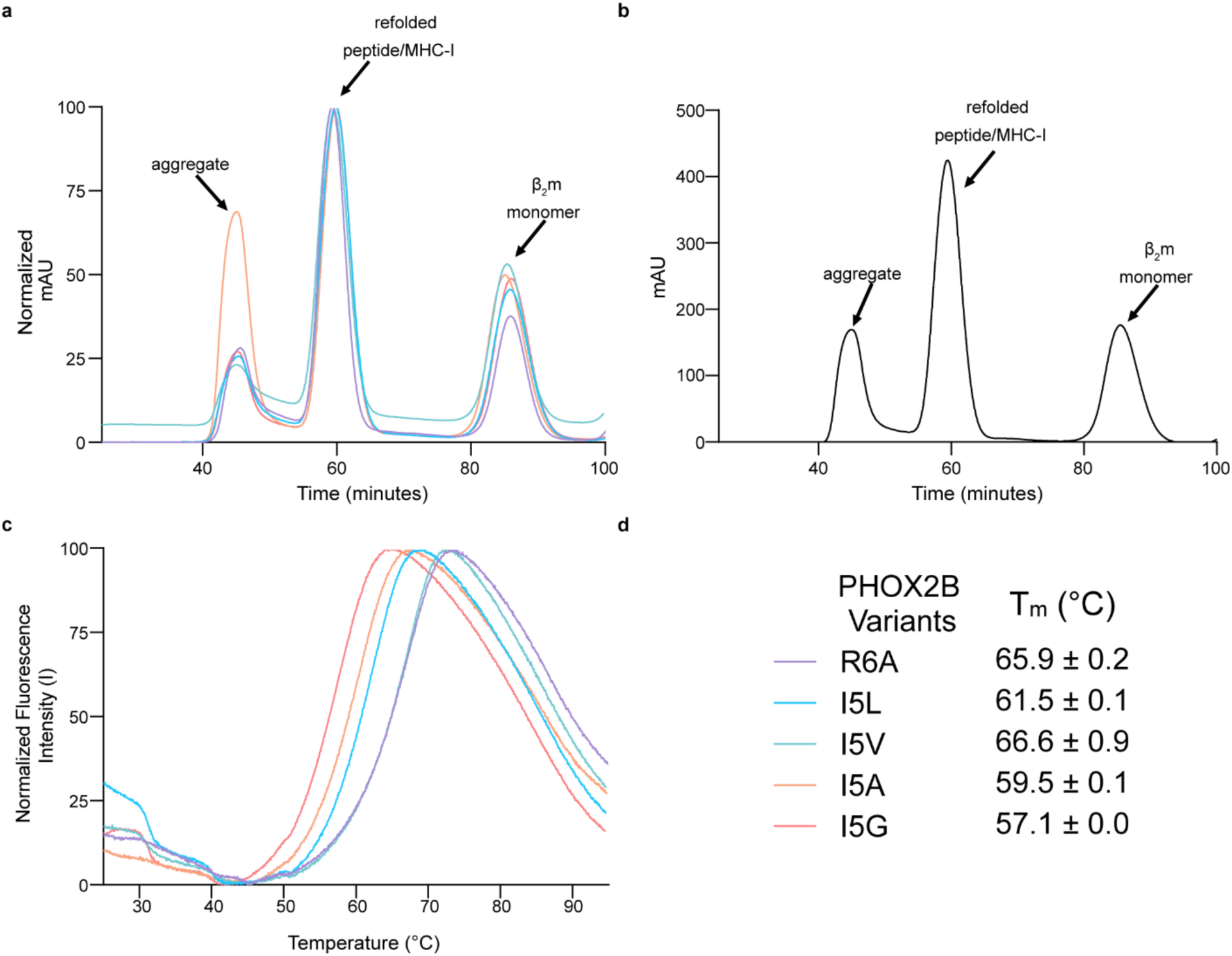
SEC and DSF data for I5 scanned PHOX2B peptides. **a**, Normalized size exclusion chromatography (SEC) traces of HLA-A*24:02/β_2_m refolded with I5 scanned PHOX2B peptides. SEC traces for predicted HLA allotypes molecules are color-coded, as shown in panel **d**. **b,** Representative SEC trace of PHOX2B/HLA-A*24:02/β_2_m indicating each peak identify. **c,** Normalized DSF traces of HLA-A*24:02/β_2_m refolded with I5 scanned PHOX2B peptides, color-coded as shown in panel **d**. **d,** Summary of melting temperatures (T_m_, °C) obtained from DSF experiments.

**Supplementary Figure 8.**
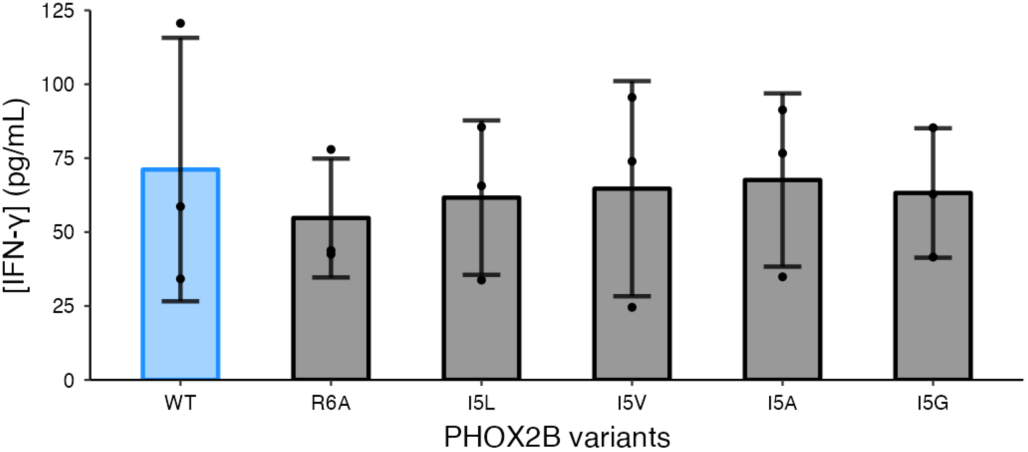
I5 scanning PHOX2B peptide cytotoxicity controls. IFN-γ levels in supernatant of mock T cells (non-transduced) co-cultured with HLA-A*24:02 colorectal adenocarcinoma cell line SW620 pulsed with PHOX2B and PHOX2B positional mutants. Exogenous peptide was added to SW620 target cells (15 µM final concentration). After four hours of incubation, T cells were added to target cells at a 3:1 ET ratio. Supernatant was collected 24 hours post effector cell addition and IFN-γ levels were measured by ELISA. Values represent mean ± SD using effector cells from n = 3 biological donors matched to CAR-T cell donors, in triplicate.

**Supplementary Figure 9.**
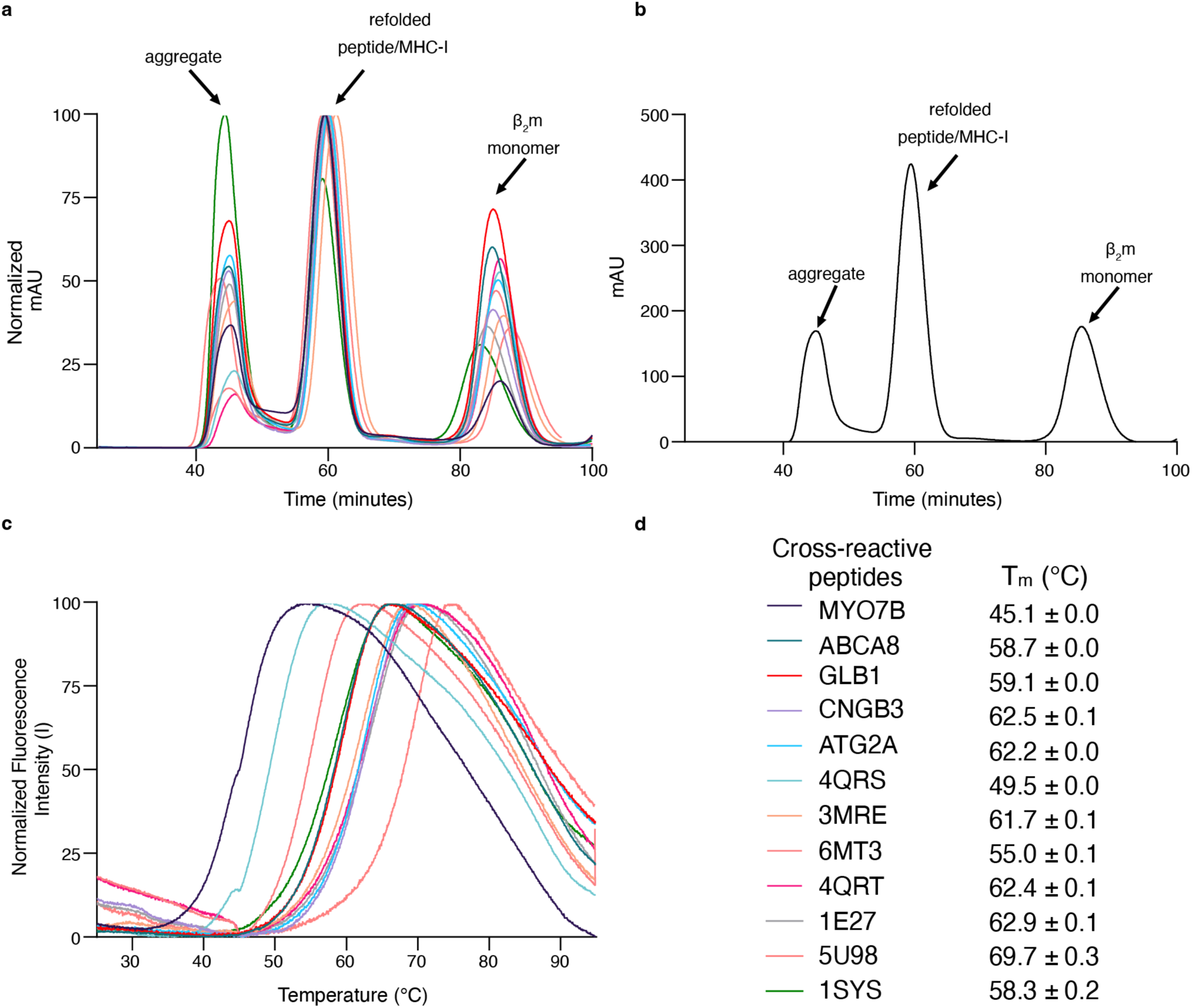
SEC and DSF data for cross-reactive peptides. **a**, Normalized size exclusion chromatography (SEC) traces of HLA-A*24:02/β_2_m refolded with cross-reactive peptides. SEC traces for HLA-A*24:02/β_2_m molecules refolded with cross-reactive peptides are color-coded, as shown in panel **d**. **b,** Representative SEC trace of PHOX2B/HLA-A*24:02/β_2_m indicating each peak identify. **c,** Normalized DSF traces of HLA-A*24:02/β_2_m refolded with cross-reactive peptides, color-coded as shown in panel **d**. **d,** Summary of melting temperatures (T_m_, °C) obtained from DSF experiments.

**Supplementary Figure 10.**
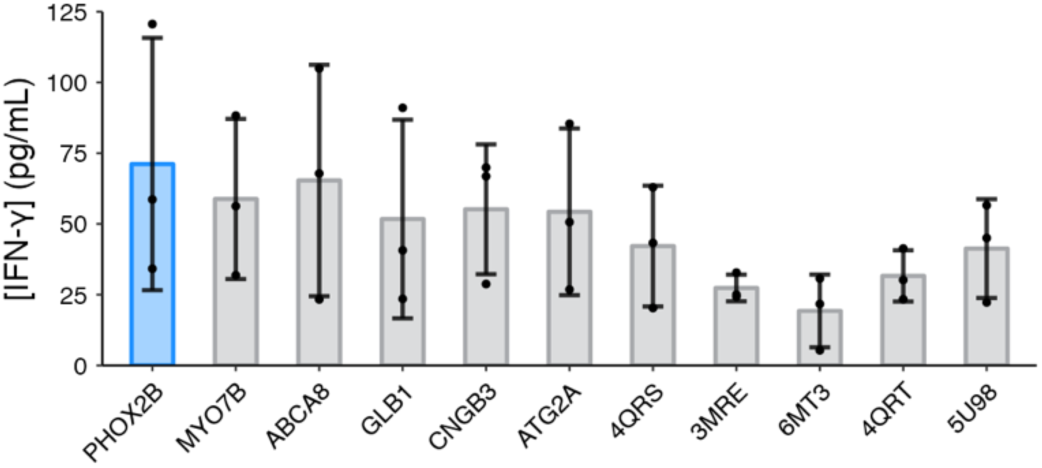
Cross-reactive peptides cytotoxicity controls. IFN-γ levels in supernatant of mock T cells (non-transduced) co-cultured with HLA-A*24:02 colorectal adenocarcinoma cell line SW620 pulsed with PHOX2B and cross-reactive peptides. Exogenous peptide was added to SW620 target cells (15 µM final concentration). After four hours of incubation, T cells were added to target cells at a 3:1 ET ratio. Supernatant was collected 24 hours post effector cell addition and IFN-γ levels were measured by ELISA. Values represent mean ± SD using effector cells from n = 3 biological donors matched to CAR-T cell donors, in triplicate.

**Supplementary Table 1.**
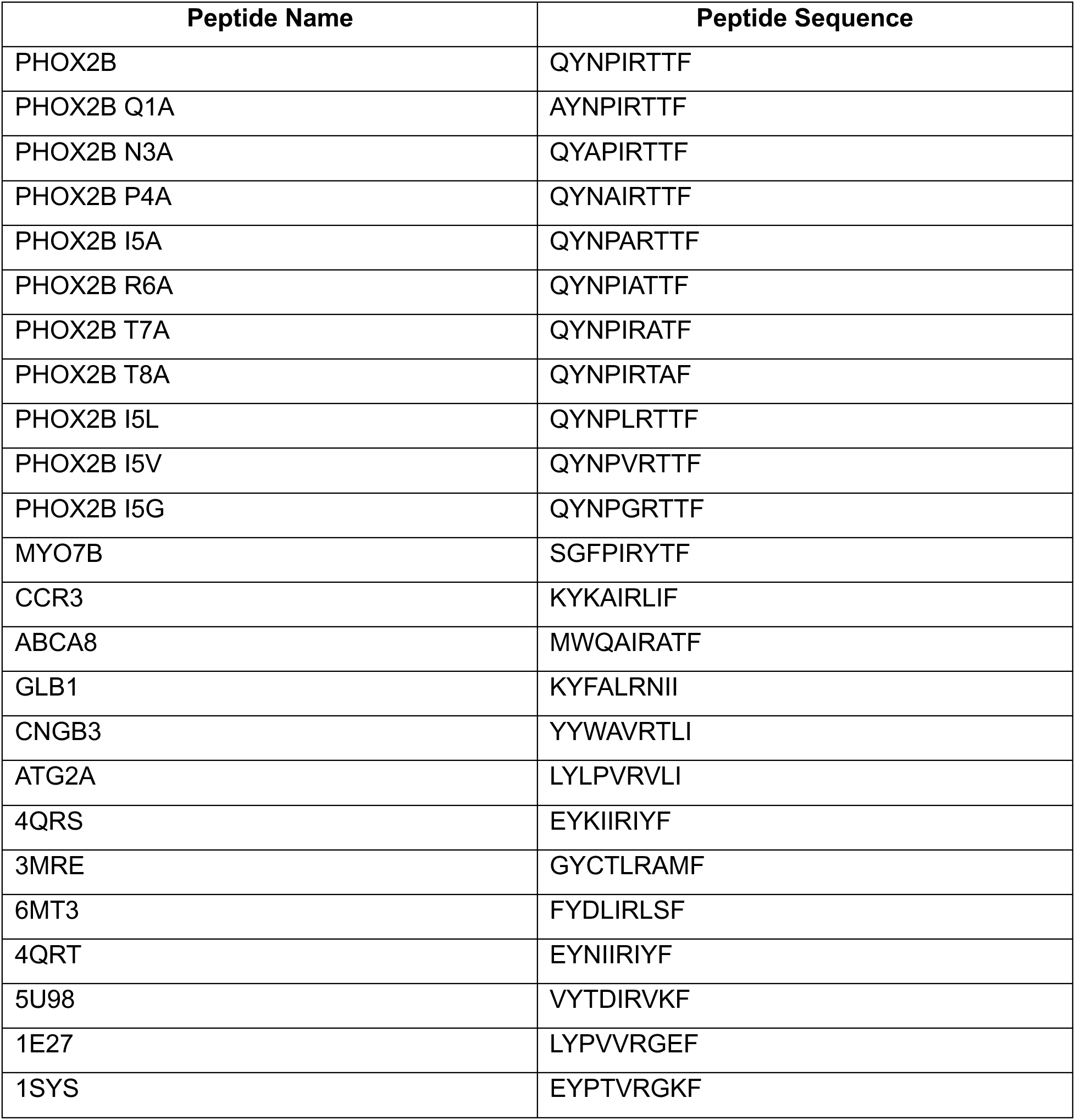
Table summarizing peptides used in this study.

**Supplementary Video 1. Interactions in the 10LH:PHOX2B/HLA-A*24:02 complex**

**Supplementary Video 2. Comparison of the crystal structure of the 10LH:PHOX2B/HLA-A*24:02 complex to the corresponding AlphaFold model.**

